# HIPK4 is essential for murine spermiogenesis

**DOI:** 10.1101/703637

**Authors:** J. Aaron Crapster, Paul G. Rack, Zane J. Hellmann, Joshua E. Elias, John J. Perrino, Barry Behr, Yanfeng Li, Jennifer Lin, Hong Zeng, James K. Chen

## Abstract

Mammalian spermiogenesis is a remarkable cellular transformation, during which round spermatids elongate into chromatin-condensed spermatozoa. The signaling pathways that coordinate this process are not well understood, and we demonstrate here that homeodomain-interacting protein kinase 4 (HIPK4) is essential for spermiogenesis and male fertility in mice. HIPK4 is predominantly expressed in round and early elongating spermatids, and *Hipk4* knockout males are sterile, exhibiting phenotypes consistent with oligoasthenoteratozoospermia. *Hipk4* mutant sperm have reduced oocyte binding and are incompetent for *in vitro* fertilization, but they can still produce viable offspring via intracytoplasmic sperm injection. Ultrastructural analyses of HIPK4-null male germ cells reveal defects in the filamentous actin (F-actin)-scaffolded acroplaxome during spermatid elongation and abnormal head morphologies in mature spermatozoa. We further observe that HIPK4 overexpression induces branched F-actin structures in cultured fibroblasts, supporting a role for this kinase in cytoskeleton remodeling. Our findings establish HIPK4 as an essential regulator of sperm head shaping and potential target for male contraception.

## INTRODUCTION

Spermiogenesis is a critical, post-meiotic phase of male gametogenesis defined by the differentiation of spermatids into spermatozoa (Figure 1A-B) (*Russell, et al., 1990*). This dramatic morphological transformation is mediated by a series of cytological processes that are unique to the testis (*Kierszenbaum, et al., 2007*; *O’Donnell, 2014*). In round spermatids, Golgi-derived vesicles give rise to the acrosome (*Berruti, et al., 2011*), a cap-like structure that is anchored to the anterior nuclear membrane by a filamentous actin (F-actin)- and keratin-containing plate called the acroplaxome (*Kierszenbaum, et al., 2003*; *Kierszenbaum, et al., 2004*). Neighboring Sertoli cells form an apical specialization that circumscribes each spermatid head, and F-actin hoops within these anchoring junctions apply external forces to the acrosome-acroplaxome complex and underlying spermatid nucleus (*Wong, et al., 2008*). The posterior nuclear pole in spermatids is simultaneously girdled by a transient microtubule- and F-actin-scaffolded structure called the manchette, which extends from the basal body of the developing flagellum and is separated from the acrosome-acroplaxome complex by a narrow groove (*Lehti, et al., 2016*).

**Figure 1.**
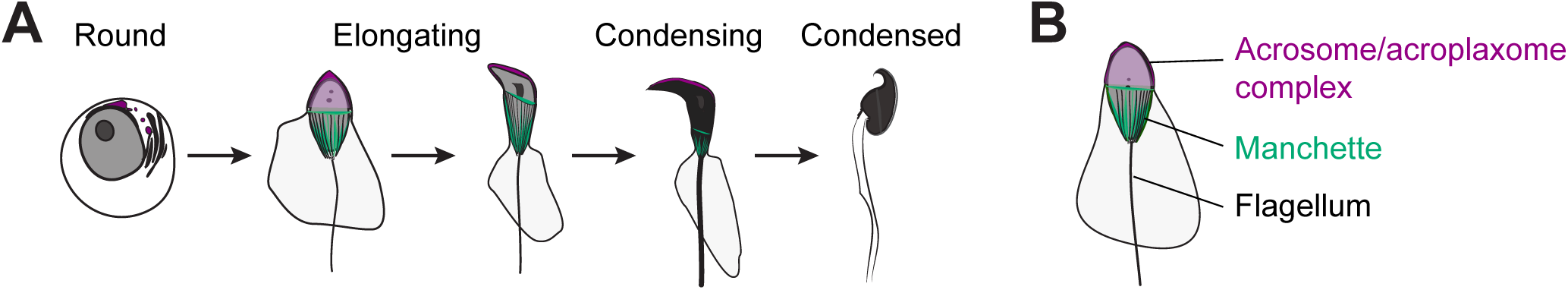
Key steps of spermiogenesis. (A) Schematic representation of murine male germ cells transitioning from round spermatids to condensed spermatozoa. These steps occur within the testis seminiferous epithelium and are conserved in all mammals. (B) Illustration of an elongating spermatid highlighting structural features that are established during spermiogenesis.

As spermiogenesis proceeds, these membranous and cytoskeletal structures act in concert to elongate the spermatid head. The spermatid nucleus becomes highly compact as chromatin condenses into a quiescent state (*Rathke, et al., 2014*), and cytoplasmic contents of the germ cell are also expelled through residual bodies or intercellular bridges to Sertoli cells (*Sprando, et al., 1987*; *Zheng, et al., 2007*). The acrosome-acroplaxome complex and manchette concurrently mediate protein transport from the Golgi to the developing flagellum, delivering cargoes required for flagellum assembly and function(*Kierszenbaum, et al., 2011*; *Kierszenbaum, et al., 2004*). In mature sperm, the acrosome then promotes sperm-egg fusion through the exocytotic release of digestive enzymes and the display of oocyte-binding receptors that are localized to the inner acrosomal membrane (*Stival, et al., 2016*).

In contrast to these detailed cytological descriptions, our understanding of the molecular mechanisms that coordinate spermiogenesis is still nascent. Initial insights into this process have been provided by mouse mutants with spermatogenic and male fertility defects (*de Boer, et al., 2015*; *Yan, 2009*). For example, pro-acrosomal vesicles fail to fuse in mice that lack the nucleoporin-like protein HRB/AGFG1 (*Kang-Decker, et al., 2001*; *Kierszenbaum, et al., 2004*), nuclear membrane protein DPY19L2 (*Pierre, et al., 2012*), certain Golgi-associated proteins [GOPC (*Yao, et al., 2002*), PICK1 (*Xiao, et al., 2009*), and GM130 (*Han, et al., 2017*)], or acrosomal factors [SPACA1 (*Fujihara, et al., 2012*) and SPATA16 (*Fujihara, et al., 2017*)]. These mutants consequently produce acrosome-less sperm with rounded heads—defects that are characteristic of globozoospermia. Acrosome biogenesis also requires the matrix component ACRBP (*Kanemori, et al., 2016*) and the coiled coil protein CCDC62 (*Li, Y., et al., 2017*; *Pasek, et al., 2016*), and loss of either acrosomal protein can cause phenotypes resembling oligoasthenoteratozoospermia (OAT), a fertility disorder characterized by low sperm concentrations and spermatozoa with abnormal shapes and reduced motility (*Tuttelmann, et al., 2018*).

Murine models have similarly revealed proteins that are required for manchette and flagellum assembly, including the RIMBP3-HOOK1 (*Zhou, et al., 2009*), LRGUK1-HOOK2 (*Liu, et al., 2015*), MEIG1-PACRG-SPAG16L (*Li, W., et al., 2015*), and FU (*Nozawa, et al., 2014*). Manchette shaping and degradation are also essential for sperm development, as demonstrated by the OAT-like phenotypes of mice expressing a loss-of-function variant of the microtubule-severing protein Katanin p80 (*O’Donnell, et al., 2012*). As spermiogenesis proceeds, membranous and cytoskeletal structures are dynamically coupled by distinct LINC (Linker of Nucleoskeleton and Cytoskeleton) complexes. These include LINC components that reside in the outer acrosomal membrane (SUN1 and nesprin3) (*Gob, et al., 2010*) or posterior nuclear envelope (SUN3, SUN4, SUN5, and nesprin1) (*Gob, et al., 2010*; *Pasch, et al., 2015*; *Shang, Y., et al., 2017*). For example, loss of SUN4 function in mice causes manchette disorganization, sperm head defects, and male sterility.

Factors that specifically contribute to acroplaxome function have been more difficult to identify and study. Actin-binding proteins such as myosins Va and VI, profilins III and IV, and cortactin localize to the acroplaxome and have been implicated in its regulation (*Behnen, et al., 2009*; *Kierszenbaum, et al., 2003*; *Kierszenbaum, et al., 2008*; *Kierszenbaum, et al., 2011*; *Zakrzewski, et al., 2017*); however, their wide-spread expression in somatic tissues has hindered functional studies. One notable exception is the actin capping protein, CAPZA3, a spermatid-specific factor that associates with CAPZB3 and binds to the barbed ends of F-actin. *Capza3* mutant male mice are sterile and have OAT-like defects, indicating that F-actin dynamics within the acroplaxome play an important role in spermiogenesis (*Geyer, et al., 2009*).

Upstream signaling proteins that control cytoskeletal dynamics are likely to be critical drivers of spermatid differentiation. For instance, PLCγ-1 phosphorylation is dysregulated in the germ cells of KIT^D814Y^ mutant mice, leading to mislocalized manchettes and deformed spermatid heads (*Schnabel, et al., 2005*). Phosphoproteomic analyses indicate that several kinase-dependent pathways are active throughout sperm development, but the roles of specific kinases in spermiogenesis are not well understood (*Castillo, et al., 2019*). Here we describe an essential function for homeodomain-interacting protein kinase 4 (HIPK4) in murine spermiogenesis and fertility. This dual-specificity kinase is predominantly expressed in the testis, where it is restricted to round and early elongating spermatids. Male *Hipk4* knockout mice are sterile and exhibit spermatogenic defects characteristic of OAT. Sperm produced by these mutant mice are also incompetent for oocyte binding and *in vitro* fertilization, and they exhibit head defects associated with dysregulation of the acrosome-acroplaxome complex. Consistent with these observations, HIPK4 overexpression in cultured somatic cells remodels the F-actin cytoskeleton and alters the phosphorylation state of multiple actin-interacting proteins. Taken together, our studies demonstrate that HIPK4 regulates acrosome-acroplaxome dynamics, spermatid head shaping, and ultimately, sperm function.

## RESULTS

### HIPK4 is expressed in differentiating spermatids

Gene expression data available through the Genotype Tissue Expression Project (https://www.gtexportal.org) and the Mammalian Reproductive Genetics Database (http://mrgd.org) indicate that HIPK4 is largely expressed in the testis, with lower levels detected in the brain. Using a tissue cDNA array and quantitative PCR, we also found that *Hipk4* is robustly transcribed in the adult murine testis (Figure 2A). *In situ* hybridization of testis sections obtained from 8-week-old C57BL/6NJ mice revealed that *Hipk4* is transcribed specifically in round and early elongating spermatids (Figure 2B), and we observed comparable *HIPK4* expression patterns in adult human testis samples (Figure 2C). We then assayed testis sections from juvenile male mice of different ages to determine precisely when *Hipk4* is expressed during spermatogenesis, taking advantage of the initial, synchronized wave of male germ cell development. *Hipk4* transcripts were first detected in germ cells at 21 days postpartum (dpp), coinciding with the appearance of step 2-3 round spermatids (Figure 2 – figure supplement 1). The population of *Hipk4*-positive spermatids expanded until 29 dpp, at which point *Hipk4* mRNA became undetectable in elongating spermatids circumscribing the seminiferous lumen. These results suggest that HIPK4 specifically functions within male germ cells as they transition from round to elongating spermatids.

**Figure 2.**
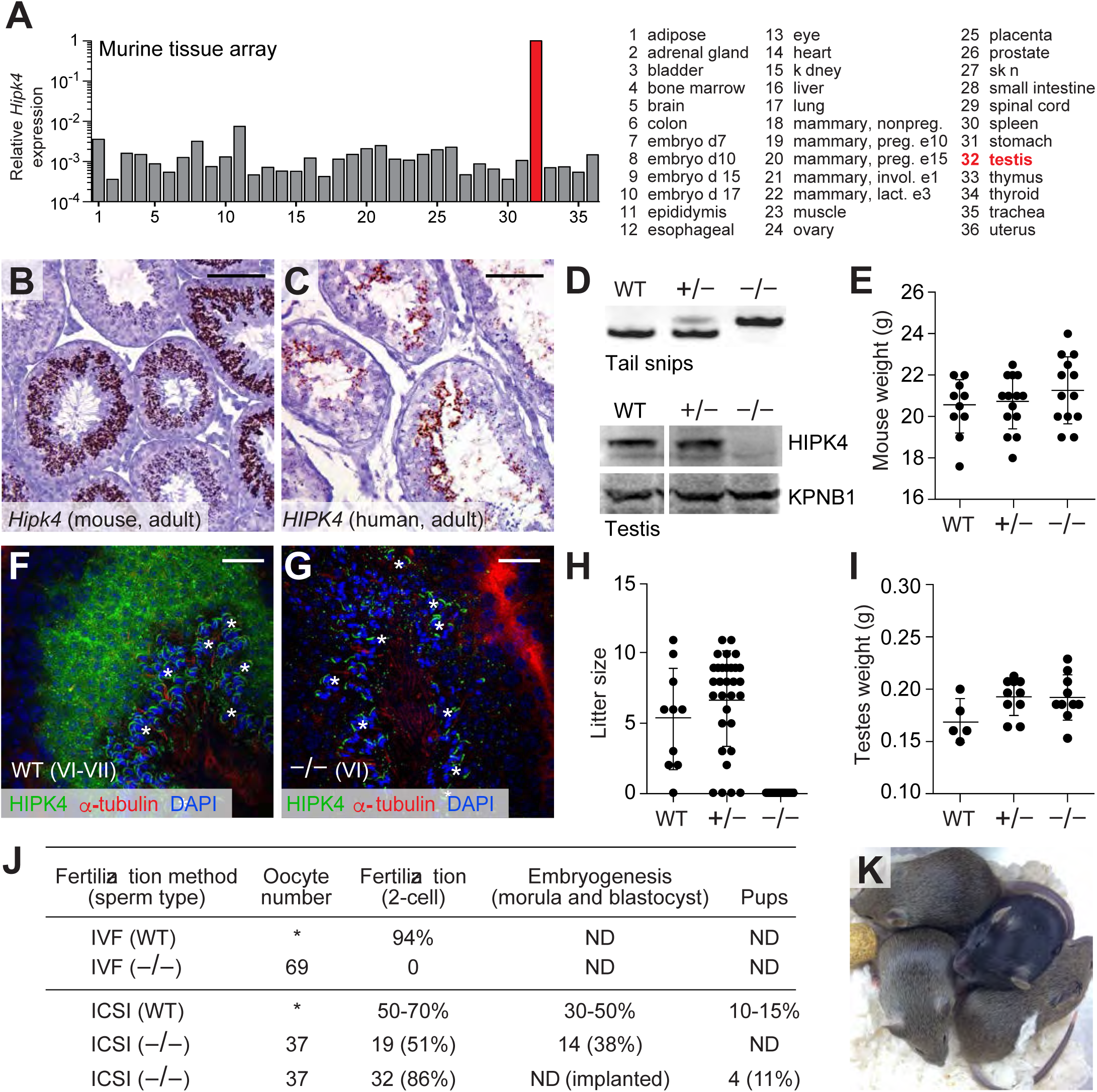
HIPK4 is expressed in spermatids and required for male fertility in mice. (A) *Hipk4* expression in various murine tissues as determined by qPCR analysis of the O rigene TissueScanTM Mouse Normal cDNA array. Data are normalized to *Gapdh*. (B-C) *Hipk4* expression in adult mouse (B) and human (C) testis sections as determined by *in situ hybridization*. (D) Validation of *Hipk4* knockout by PCR of tail-derived genomic DNA and western blot analyses of whole testis lysates. Immunoblots are from the same membrane and exposure time. (E) Weights of WT, *Hipk4*^+/−^, and *Hipk4*^−/−^ males at 6-7 weeks of age. (F-G) HIPK4 protein expression in adult mouse seminiferous tubule sections (Stage VI-VII) as determined by immunofluorescence imaging. Asterisks indicate non-specific antibody binding (see Figure 2 − figure supplement 2). (H) Number of live pups per litter resulting from crosses between 7-week-old males and age-matched, WT females. (I) WT, *Hipk4*^+/−^ and *Hipk4*^−/−^ testis weights at 6 weeks of age. (J) Fertilization potential of *Hipk4*^−/−^ sperm using IVF and ICSI. ND = not determined. Experiments were conducted by the Stanford Transgenic, Knockout, and Tumor Model Center, and the wild-type data represents the core facility’s average results using C57 BL/6 NJ sperm. (K) Pups born via ICSI using *Hipk4*^−/−^ sperm. Scale bars: B-C, 10 0 μm; F-G, 20 μm. Statistical analyses: error bars depicted in panels E, H, and I represent the average value ± s.d.

### HIPK4 is essential for male fertility

*Hipk4* knockout mice were first reported in a 2008 patent application by Bayer Schering Pharma, which described general defects in sperm morphology and number (*Sacher, et al., 2008*). However, the fertility of these mutant mice was not characterized, nor were the mice made publicly available. As part of the Knockout Mouse Phenotyping Program (KOMP^2^), the Jackson Laboratory generated mice containing a *Hipk4* null allele (*tm1b*), in which a β-galactosidase reporter replaces exons 2 and 3. We established a colony of *Hipk4^tm1b/tm1b^* mice (henceforth referred to as *Hipk4*^−/−^) and confirmed that these mice fail to produce functional *Hipk4* gene products using *in situ* hybridization and western blot analysis (Figure 2D). Loss of HIPK4 had no apparent effect on the animal viability or growth (Figure 2E). By immunostaining testis cryosections from adult wild-type and *Hipk4*^−/−^ mice, we confirmed that HIPK4 protein is expressed in round and early elongating spermatids (steps 3-8). The kinase is distributed throughout the cytoplasm of these germ cells, mirroring its subcellular localization when overexpressed in somatic cells (Figure 2F-G, Figure 2 – figure supplement 2) (*van der Laden, et al., 2015*).

Despite their grossly normal development and physiology, homozygous *Hipk4* mutant males were unable to conceive, whereas heterozygote males sired normal litter sizes (Figure 1H-I). No significant differences in testis weight were observed across the *Hipk4* genotypes (Figure 1J). Female *Hipk4*^−/−^ mice were physically indistinguishable from their wild-type and heterozygous littermates and gave birth to normal litter sizes (data not shown). Epididymal sperm isolated from *Hipk4*^−/−^ mice failed to fertilize wild-type oocytes under standard *in vitro* fertilization (IVF) conditions, but intracytoplasmic sperm injection (ICSI) of the mutant sperm yielded embryos that could undergo uterine implantation to produce healthy pups (Figure 2J-K).

### HIPK4-deficient mice exhibit OAT-like phenotypes

During our *in vitro* fertilization studies, it was apparent that the *Hipk4*^−/−^ male mice had spermatogenesis defects consistent with OAT. In comparison to wild-type mice, homozygous mutants produced sperm at low epididymal concentrations, and the germ cells had decreased motility [both the total number of motile sperm and those with progressive motility as measured by computer-assisted sperm analysis (CASA)] and abnormal morphology (Figure 3A-E, Figure 3 – figure supplement 1). Head defects included macrocephaly, microcephaly, and irregular shapes; tail deformities included bent, coiled, crinkled, and shortened flagella. *Hipk4^+/–^* sperm also had reduced epididymal concentrations and total motility, but their progressive motility and morphology were normal. We observed that over 10% of the epididymal sperm isolated from *Hipk4*^−/−^ mice exhibited DNA fragmentation that could be detected by TUNEL (terminal deoxynucleotidyl transferase dUTP nick end labeling) staining. In comparison, only 1.5% of wild-type or *Hipk4*^+/–^ sperm were TUNEL-positive. Homozygous mutant sperm therefore may undergo a higher rate of apoptosis, possibly accounting for their lower epididymal concentrations.

**Figure 3.**
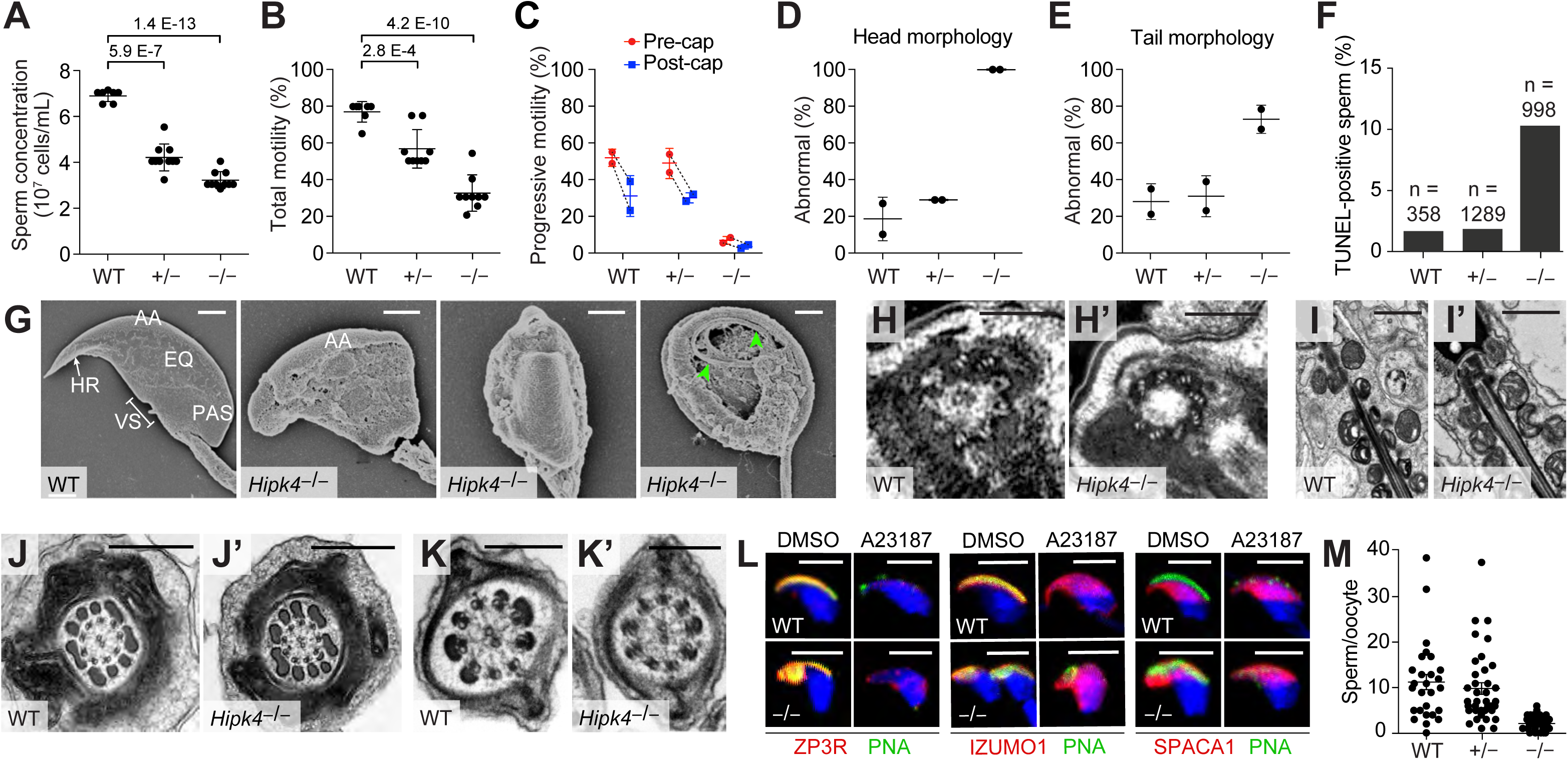
*Hipk4* knockout mice exhibit oligoasthenoteratozoospermia. (A) Concentrations of epididymal sperm ± s.d. *P* values for the indicated statistical comparison are shown. (B) Percentage of epididymal sperm that were motile ± s.d. *P* values for the indicated statistical comparison are shown. (C) CASA measurements of progressive sperm motilty before and after capacitation ± s.d. Data connected by the dashed lines are showing sperm motility from the same animal. (D, E) Percentage of sperm morphology with abnormal head or tail morphology ± s.d. as assessed by phase contrast microscopy (see also, Figure 3 − Figure supplement 1). (F) Quantification of TUNEL-positive epididymal sperm with the designated genotypes (n = total number of sperm analyzed). (G) SEM images of epididymal sperm. AA = anterior acrosome, EQ = eq uatorial segment, PAS = postacrosomal segment, VS = ventral spur, HR = hook rim. Arrowheads point to the axoneme wrapped inside a demembranated sperm head. (H, H’) Centrioles at the basal body of step 15 spermatid axonemes. (I, I’) Mitochondria along the midpiece of step 15 spermatids. (J, J’) Cross section of step 15 spermatid flagella at the midpiece. (K, K’) Cross section of step 15 spermatid flagella at their principal piece. (L) Acrosomal changes in capacitated sperm treated with a Ca^2+^ ionophore (A23187) as assessed by immunofluorescence and staining with FITC-labeled PNA. Nuclei were stained with DAPI. (M) Number of oocyte-bound sperm under standard IVF conditions after extensive washing ± s.e.m. Scale bars: G, 2 μm; H, 1 μm; I, 1 μm; J, 0.5 μm; K, 0.2 μm; L, 5 μm.

We further compared the head structures of wild-type and *Hipk4*^−/−^ sperm by scanning electron microscopy (SEM; Figure 3G). All HIPK4-deficient sperm exhibited head morphologies that deviated from the flat, crescent-shaped structures of their wild-type counterparts. Specific defects included a disorganized anterior acrosome and the absence of a distinct equatorial segment, post-acrosomal sheath, ventral spur, and sharp hook rim. In some sperm samples with demembranated head structures, we observed mislocalized axonemal components wrapped around the nucleus. We also used transmission electron microscopy (TEM) to analyze the tail structures of newly formed spermatozoa within the seminiferous tubule. HIPK4 loss did not appear to affect the basal body (Figure 3H-H’), axoneme (Figure 4I-K’), outer dense fibers (Figure 4I-I’), mitochondria (Figure 4J-J’), or fibrous sheath (Figure 4K-K’) of these fully differentiated cells.

**Figure 4.**
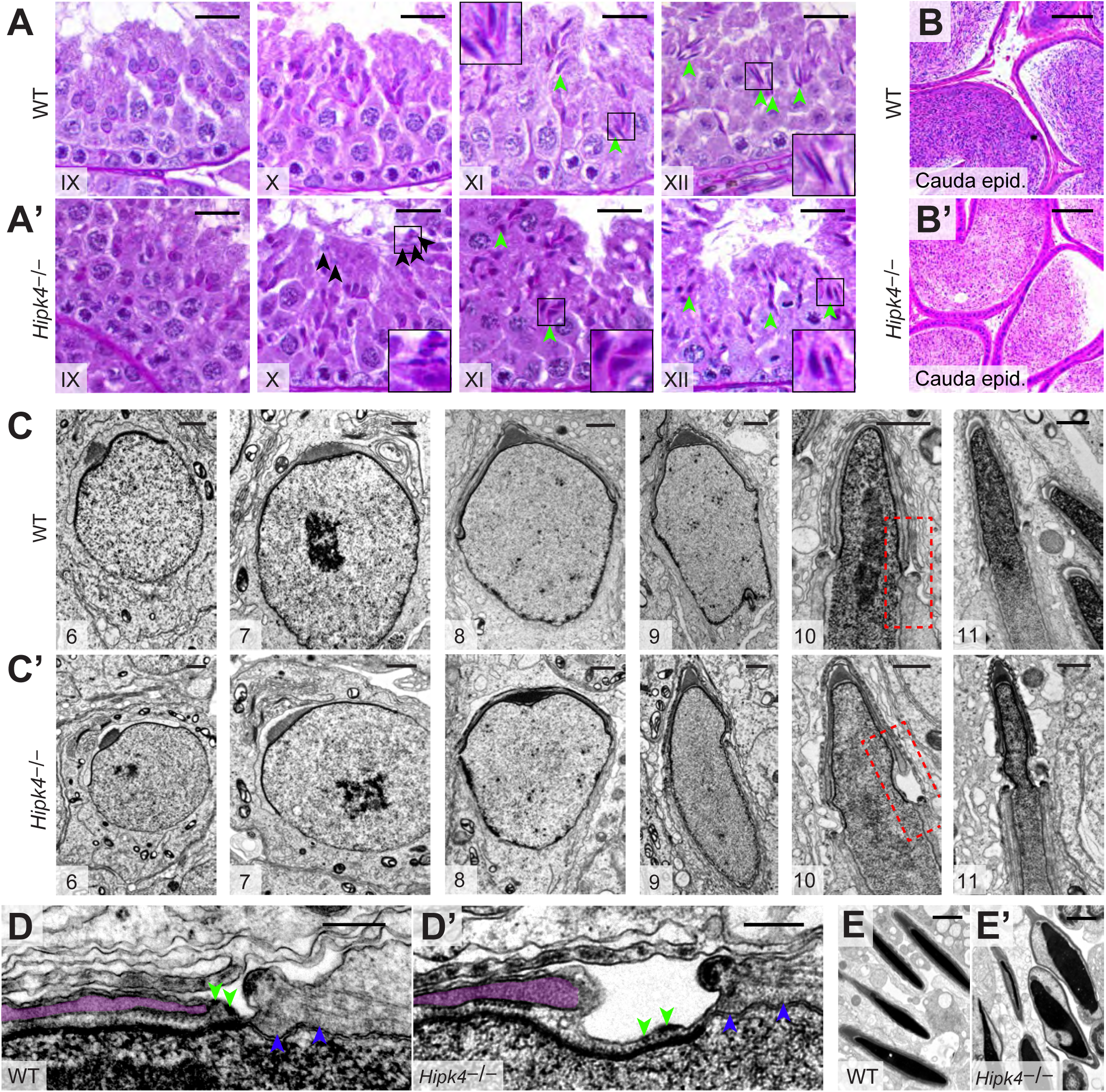
HIPK4 regulates the acrosome-acroplaxome complex. (A,A’) PAS-stained sections of seminiferous tubules from WT or *Hipk4*^−/−^ mice at various stages of spermatogenesis. Black arrowheads point to testicular spermatozo a that are retained in a stage X *Hipk4*^−/−^ tubule. G reen arrowheads point to the heads of elongating spermatids. (B, B’) H& E-stained sections of caudal epididymides from WT or *Hipk4*^−/−^ mice. (C-E) TEM images. (C, C’) Step 6-11 spermatids from WT or *Hipk4*^−/−^ mice. Areas outlined by the red dashed boxes are shown at higher magnification in D-D’. (D, D’) Magnified view of the groove belt region of step 10 spermatids. G reen arrowheads point to electron densities in the acroplaxome that are normally associated with the posterior edge of the acrosome (highlighted in purple). Blue arrowheads point to filaments linki ng the manchette to the nuclear envelope. (E, E’) TEM images of condensed spermatids in WT and *Hipk4*^−/−^ testis sections. Scale bars: A-B, 20 μm; C, 1 μm; D, 0.2 μm; E, 2 μm.

### HIPK4-deficient sperm can undergo capacitation and the acrosome reaction in vitro

We next examined how the loss of HIPK4 affects two key aspects of sperm function: capacitation and the acrosome reaction. Sperm naturally undergo capacitation as they ascend the female reproductive tract and interact with the oviduct epithelium (*Austin, 1951*; *Chang, 1951*). Changes in the glycoproteins, phospholipids, and cholesterol residing in the sperm plasma membrane, Ca^2+^ influx, and increases in intracellular pH correlate with the activation of soluble adenylate cyclase and protein kinase A (*Abou-haila, et al., 2009*; *Lin, et al., 1996*; *O’Rand, 1982*; *Stival, et al., 2016*). Downstream protein tyrosine phosphorylation signaling events are initiated, and sperm switch from progressive motility to a “hyperactivated” swimming motion (*Alvau, et al., 2016*; *Naz, et al., 2004*; *Sepideh, et al., 2009*). Capacitated sperm also become competent for the acrosome reaction, during which the outer acrosomal membrane fuses with the overlying plasma membrane (*Hirohashi, 2016*; *Kierszenbaum, 2000*; *Stival, et al., 2016*). This process causes the release of digestive enzymes stored within the acrosome, and it exposes oocyte-binding receptors that are displayed on the inner acrosomal membrane, which spreads down over the equatorial region of the sperm head (*Sebkova, et al., 2014*; *Sosnik, et al., 2009*). Together, these steps ultimately promote sperm-oocyte engagement, fusion, and fertilization.

To determine whether HIPK4 is required for sperm capacitation, we isolated motile sperm from the caudal epididymides of wild-type, *Hipk4*^+/–^, and *Hipk4*^−/−^ mice and incubated them in capacitation medium containing Ca^2+^, bicarbonate, and bovine serum albumin. We then assessed the resulting levels of tyrosine phosphorylation (p-Tyr) in soluble sperm lysates by western blot. We observed no significant differences in p-Tyr between the three *Hipk4* genotypes (Figure 3 – figure supplement 2A), and by CASA, we found that wild-type, *Hipk4*^+/–^, and *Hipk4*^−/−^ sperm treated with capacitation medium undergo similar changes in progressive motility (Figure 3C). These results indicate that *Hipk4*^−/−^ sperm are competent for capacitation *in vitro*.

To investigate the ability of HIPK4-deficient sperm to undergo acrosomal exocytosis, we incubated capacitated sperm with the Ca^2+^ ionophore A23187. Sperm were then fixed, quickly permeabilized, and labeled with fluorescein-conjugated peanut agglutinin (FITC-PNA), which binds to the outer acrosome membrane and is lost upon exocytosis. After 1.5 hours of capacitation, 86% of wild-type sperm had fully intact acrosomes, whereas only 63% of mutant sperm retained acrosomes (due to either head malformations or spontaneous acrosome exocytosis). As expected, A23187 treatment decreased the percentage of FITC-PNA positive cells in both wild-type and *Hipk4*^−/−^ samples (to 22% and 45%, respectively), although mutant sperm responded less efficiently to this treatment.

To further assess the acrosome reaction in *Hipk4*^−/−^ sperm, we examined specific acrosomal proteins during A23187-induced exocytosis: ZP3R, IZUMO1, and SPACA1. ZP3R (also known as sp56) is a zona pellucida-binding protein that appears on the sperm surface after capacitation, and it is released when the outer acrosomal and plasma membranes fuse (*Kim, K. S., et al., 2001*). IZUMO1 (*Inoue, et al., 2005*; *Sebkova, et al., 2014*) and SPACA1 (*Fujihara, et al., 2012*) are membrane proteins required for head shaping and oocyte fusion that localize to distinct acrosomal regions. IZUMO1 spreads from the anterior acrosomal cap to the equatorial segment during the acrosome reaction (*Inoue, et al., 2005*; *Sebkova, et al., 2014*), and SPACA1 remains localized to the equatorial region (Figure 3H) (*Fujihara, et al., 2012*). As assessed by immunofluorescence microscopy, all three acrosomal factors exhibited HIPK4-independent behaviors in response to Ca^2+^ ionophore exposure. Together, these findings suggest that HIPK4-deficient sperm are capable of normal acrosomal changes, at least in response to calcium signaling.

### HIPK4-deficient sperm exhibit diminished oocyte binding

Since *Hipk4*^−/−^ sperm retain their ability to undergo capacitation and the acrosome reaction *in vitro*, we considered whether the head defects caused by HIPK4 loss might compromise sperm-oocyte interactions. Equivalent numbers of motile, capacitated sperm from wild-type, *Hipk4*^+/–^, or *Hipk4*^−/−^ males were incubated with cumulus-oocyte complexes (COCs) in human tubal fluid (HTF) supplemented with Ca^2+^ and glutathione. The complexes were then washed repeatedly, fixed, and oocyte-bound sperm were quantified by nuclear staining and confocal imaging (Figure 3I). Consistent with the incompetence of *Hipk4*^−/−^ sperm for IVF, these cells bound less efficiently to oocytes in comparison to their wild-type and heterozygous mutant counterparts. We further noted that the COCs incubated with mutant sperm retained many cumulus cells, while all of the cumulus cells of the COCs exposed to wild-type sperm were detached (Figure 3 – figure supplement 2). Thus, HIPK4-deficient sperm lack structural and/or molecular features that are required for maximally productive sperm-oocyte interactions.

### Loss of HIPK4 function disrupts the acrosome-acroplaxome complex

To gain insights into the spermatogenic defects caused by loss of HIPK4, we compared periodic acid-Schiff (PAS)-stained testis sections obtained from adult wild-type and *Hipk4*^−/−^ mice. *Hipk4*^−/−^ testes contained malformed elongating spermatids, which failed to properly extend by step 12 (Figure 4A-A’, Figure 4 – figure supplement 1A-B). In addition, stage IX and X tubules from mutant mice contained spermatozoa that were still attached to Sertoli cells. Since these mature germ cells are normally released into the seminiferous lumen by stage VIII, HIPK4-deficient spermatozoa appear to have delayed spermiation, which may also contribute to the lower sperm counts observed in *Hipk4*^−/−^ mice. Despite their morphological abnormalities, *Hipk4*^−/−^ spermatozoa released to the epididymis progressed normally to the cauda (Figure 4B-B’, Figure 4 – figure supplement 1C-D).

Next, we analyzed spermatid structures in greater detail by TEM imaging of testis sections. Although HIPK4 protein expression peaks in step 5-7 spermatids, all *Hipk4*^−/−^ round spermatids appeared normal by TEM (Figure 4C-C’, Figure 4 – figure supplement 2A). Spermatids begin to elongate at step 8, and most of these cells appeared normal in *Hipk4*^−/−^ testes. However, some HIPK4-null step 8 spermatids contained highly amorphous, fragmented acrosomal vesicles and/or detached acrosomal granules, which were coincident with structural abnormalities to the anterior nuclear pole (Figure 4 – figure supplement 2A). Aberrant head structures became universally apparent in step 9-10 *Hipk4*^−/−^ spermatids (Figure 4C-C’). The acrosome-acroplaxome marginal ring was no longer juxtaposed to the perinuclear ring of the manchette, significantly widening the groove belt and deforming the underlying nuclear lamina (Figure 4D-D’). In some cases, enlargement of the groove belt was accompanied by detachment of the acrosome, severe anterior head deformities, and retention of spermatid cytoplasm (Figure 4 – figure supplement 2B). We also observed small electron-dense areas at the posterior boundary of the acroplaxome that have not been previously defined. These electron-dense structures appeared to interact with the acrosome in wild-type spermatids, but they were disorganized and uncoupled from this organelle in HIPK4-null cells (Figure 4D-D’, Figure 4 – figure supplement 2C).

In contrast to the acroplaxome-acrosome defects, other anterior head structures appeared to form properly in HIPK4-deficient spermatids. For example, the anterior nuclear membrane (nuclear dense lamina) is putatively anchored to the acroplaxome *via* the inner membrane protein DPY19L2, while the outer acrosomal membrane is connected to the cytoskeleton via a LINC complex composed of SUN1 and nesprin3. These membrane-cytoskeleton linkages had comparable subcellular distributions in wild-type and *Hipk4*^−/−^ spermatids, as assessed by immunofluorescence imaging of isolated germ cells (Figure 4 – figure supplement 3A-B). TEM analyses similarly revealed no overt defects within the manchette, including the perinuclear ring and the conical array of filaments that scaffold the posterior nuclear pole (Figures 4C-C’, 4D-D’). We further characterized manchette formation and degradation by immunostaining the microtubule end-binding protein EB3 in testis cryosections and isolated germ cells. Testis sections from *Hipk4*^−/−^ mice exhibited wild-type-like manchette dynamics (Figure 4 – figure supplement 3C). However, elongating *Hipk4*^−/−^ spermatids isolated from dissociated testis tissues occasionally exhibited abnormally shaped manchettes (Figure 4 – figure supplement 3D), suggesting that HIPK4 may also regulate certain aspects of this microtubule- and F-actin-scaffolded structure.

### HIPK4 does not primarily act through transcriptional regulation

Given the delay between the onset of HIPK4 expression in wild-type round spermatids and the emergence of HIPK4-dependent morphological phenotypes, we investigated whether HIPK4 signaling could regulate the acrosome-acroplaxome complex through transcriptional mechanisms. Other HIPK family members and related kinases (*e.g.*, DYRKs) localize to the nucleus and have established roles in transcriptional regulation (*Di Vona, et al., 2015*; *Rinaldo, et al., 2008*), and HIPK4 can phosphorylate p53 *in vitro* (*Arai, et al., 2007*). We therefore used oligonucleotide microarrays with full-genome coverage to profile transcriptional differences between adult testes isolated from wild-type and *Hipk4*^−/−^ mice. Through this approach, we identified 415 genes that were upregulated ≥2-fold in *Hipk4*^−/−^ testes compared to wild-type tissue and 709 genes that were downregulated ≥2-fold (Figure 5A and Figure 5 – Source data 1). Hierarchical clustering of the data from biological replicates revealed overlapping groupings of wild-type and *Hipk4*^−/−^ samples (Figure 5B), indicating that the transcriptional differences between the genotypes are modest. *Hipk4* itself exhibited the largest change in transcript abundance between genotypes (Figure 5C). We also noted that loss of HIPK4 did not significantly alter the mRNA levels of transcription factors that are unique to stage 5-7 seminiferous tubules (*Green, et al., 2018*). Based on these microarray data, we surmise that HIPK4 does not primarily act through transcriptional regulation.

**Figure 5.**
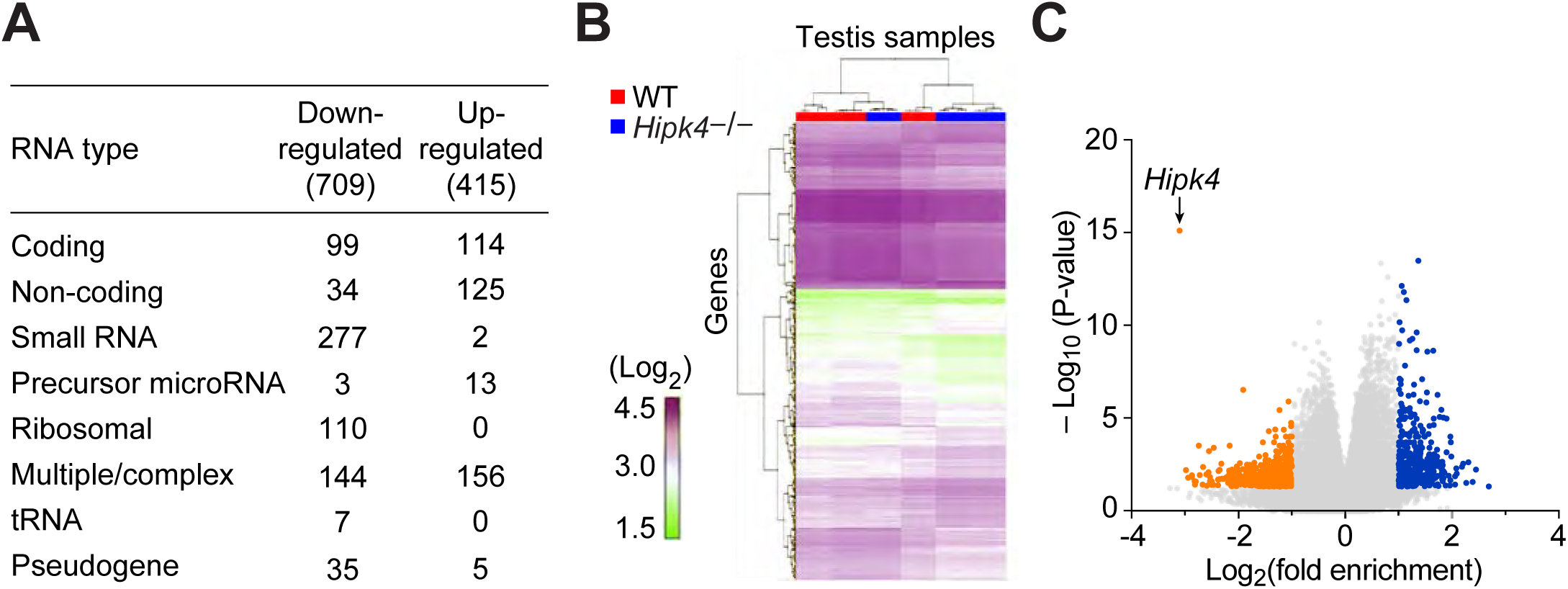
Wild-type and Hipk4 mutant testes have similar transcriptomes. (A) Summary of the types of RNA that were increased (up-regulated) or decreased (down-regulated) in *Hipk4* knockout testes compared to wild-type, as measured by quadruplicate microarray analysis of three testis samples. (B) Histogram of normalized signal intensities (Log_2_) of gene expression levels in individual microarrarys. Color scheme was arbitrarily assigned. (C) Volcano plot depicting the transcriptional differences between *Hipk4* knockout and wild-type testes. RNA species that exhibited a > 2-fold change in abundance and had a *P*-value < 0.05 are shown in orange (down-regulated) or blue (up-regulated).

### HIPK4 overexpression remodels F-actin in cultured somatic cells

In the absence of discrete HIPK4-dependent transcriptional programs, we investigated the biochemical functions of HIPK4 in cells. We retrovirally overexpressed wild-type *Hipk4* or a catalytically dead mutant, *Hipk4 K40S* (*Arai, et al., 2007*), in cultured mouse embryonic fibroblasts (NIH-3T3 cells). Untransduced fibroblasts or those expressing the K40S mutant maintained spindle-like morphologies, whereas cells expressing HIPK4 became either spherical or polygonal within two days after infection. The polygonal HIPK4-overexpressing cells were multinucleate, likely due to cytokinesis failure (Figure 6A-C). Coincident with these changes in cell shape, we observed a striking loss of F-actin-containing stress fibers in the HIPK4-overexpressing cells (Figure 6D-E). The overall ratio of globular actin (G-actin) to F-actin did not increase with *Hipk4* transduction (Figure 6F), suggesting that this kinase induces F-actin remodeling rather than filament disassembly into monomeric subunits. HIPK4 overexpression did not overtly alter the microtubule cytoskeleton in these cells (Figure 6G-H).

**Figure 6.**
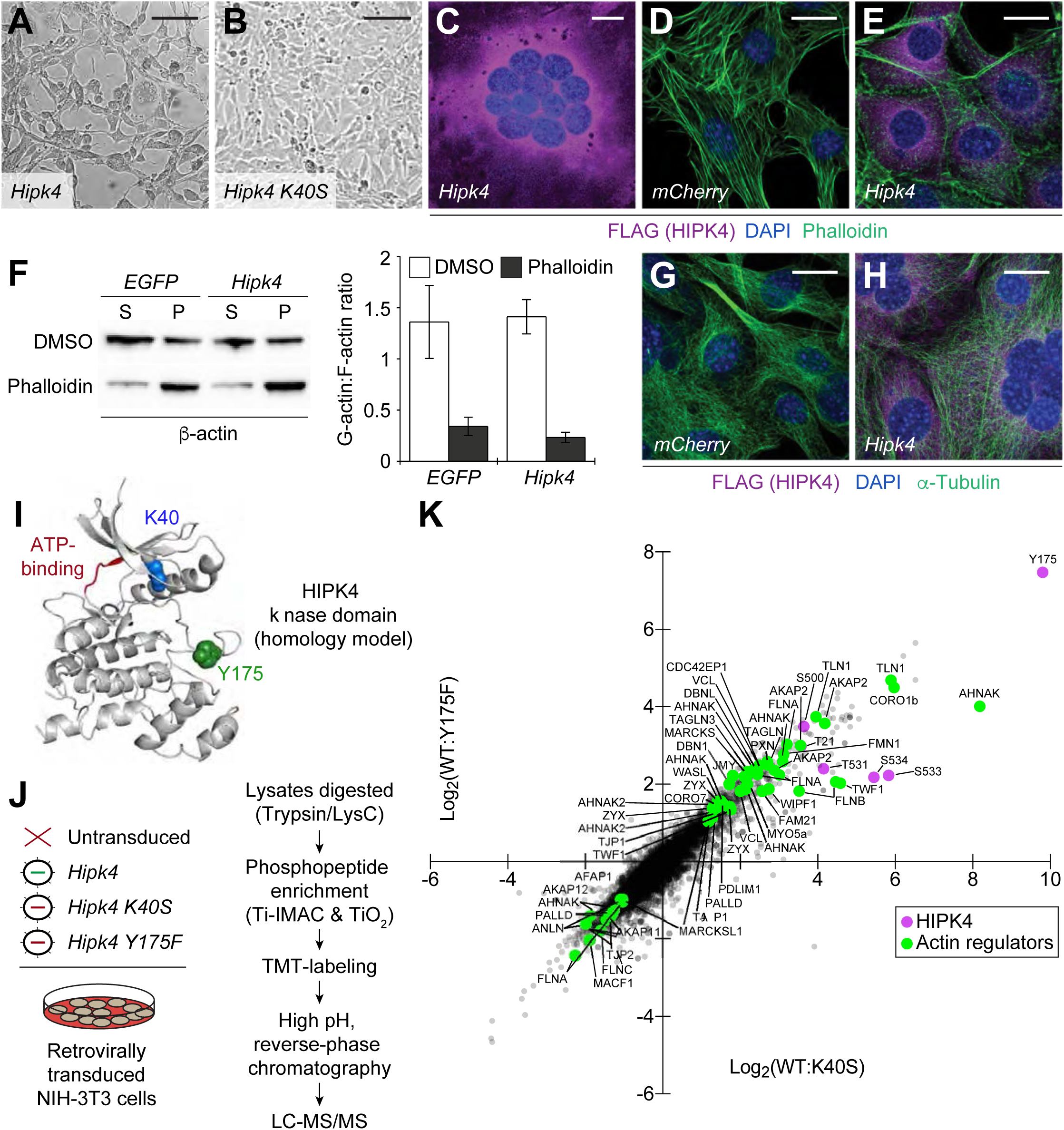
*Hipk4* overexpression remodels the actin cytoskeleton in cultured cells. (A-B) Brightfield images of NIH-3T3 cells retrovirally transduced with FLAG-tagged *Hipk4* or kinase-dead *Hipk4 K40S*. (C) HIPK4-expressing NIH-3T3 cell with multiple nuclei. (D-E) Phalloidin and anti-FLAG staining of NIH-3T3 cells transduced with *mCherry* or FLAG-tagged *Hipk4*. (F) G- and F-actin levels in NIH-3T3 cells transduced with *EGFP* or FLAG-tagged *Hipk4* and then treated with DMSO or phalloidin after lysis. The western blot shows the soluble (S; G-actin) and pelleted (P; F-actin) pools of actin after ultracentrifugation, and the graph depicts the average G-actin:F-actin ratios of quadruplicate samples ± s.e.m. (G-H) α-Tubulin and FLAG immunostaining of NIH-3T3 cells transduced with *mCherry* or FLAG-tagged *Hipk4*. (I) Homology model of the HIPK 4 ki nase domain using the DYRK 1A structure as a template (PDB ID: 3ANQ). The ATP-binding site is colored red, and residues that are essential for catalytic activity are depicted as blue or green space-filling models. (J) Work flow used to characterize the HIPK 4-dependent phosphoproteome in NIH-3T3 cells. (K) Scatter plot of 6,941 phosphosites identified by LC-MS/MS, graphed according to their relative levels in NIH-3T3 cells overexpressing wild-type or catalytically inactive HIPK 4. Selected HIPK 4-regulated phosphosites in actin-modulating proteins are shown in green and phosphosites in HIPK 4 are shown in purple. Scale bars: A-B, 100 μm; C-E and G-H, 20 μm.

We then compared the phosphoproteomes of NIH-3T3 cells transduced with wild-type HIPK4, the K40S variant, or a second inactive mutant that is incapable of autophosphorylation-dependent activation (Y175F) (*van der Laden, et al., 2015*) (Figure 6I-J). Through phosphopeptide enrichment, isobaric labeling, and quantitative mass spectrometry, we identified 6,941 phosphosites; 303 of which increased in abundance by ≥2-fold upon HIPK4 overexpression (in comparison to either inactive mutant) (Figure 6K and Figure 6 – Source data 1). Consistent with the effects of HIPK4 overexpression on F-actin dynamics, several of these phosphosites reside in known actin-interacting proteins. For example, we identified HIPK4-dependent phosphosites in talin 1 (TLN1), AHNAK nucleoprotein, coronin 1B (CORO1B), A-kinase anchor protein 2 (AKAP2), formin 1 (FMN1), vinculin (VCL), MARCKS, paxillin (PXN), WASH family members (WASL, WIPF1, and FAM21), zyxin (ZYX), unconventional myosin 5a (MYO5a), filamins (FLNA and FLNB), and transgelins (TAGLN and TAGLN3).

### Loss of HIPK4 alters F-actin dynamics in the spermatid acroplaxome

Because HIPK4 overexpression altered stress fiber dynamics in cultured cells, we hypothesized that HIPK4 may similarly modulate F-actin-related functions in developing sperm. In particular, a role for HIPK4 in regulating acroplaxome structure and/or function could explain the head defects observed in *Hipk4*^−/−^ spermatids. We first used fluorescently labeled phalloidin to visualize F-actin structures in wild-type and *Hipk4*^−/−^ testis sections; however, since the phalloidin staining predominantly labeled the F-actin hoops of Sertoli cells, it was difficult to discern differences between the samples (Figure 7A). We consequently used the phalloidin probe to analyze spermatids isolated from dissociated testes, which enabled us to specifically image F-actin in the acroplaxome. Although there was no discernable difference in acroplaxome F-actin in round-to-early elongating spermatids isolated from wild-type and *Hipk4*^−/−^ testes (Figure 7B), wild-type but not *Hipk4*^−/−^ germ cells maintained this F-actin-based plate at later stages of spermatid differentiation (e.g., condensed spermatids). To further compare F-actin-related structures in wild-type and *Hipk4*^−/−^ spermatids, we assessed the localization of actin-capping proteins CAPZA3 and CAPZB3 by immunofluorescence microscopy (Figure 7C-D). Consistent with our phalloidin staining results, elongating wild-type and *Hipk4*^−/−^ spermatids exhibited similar expression levels and subcellular distributions of CAPZA3 and CAPZB3, but these actin regulators were selectively diminished in condensing *Hipk4*^−/−^ spermatids.

**Figure 7.**
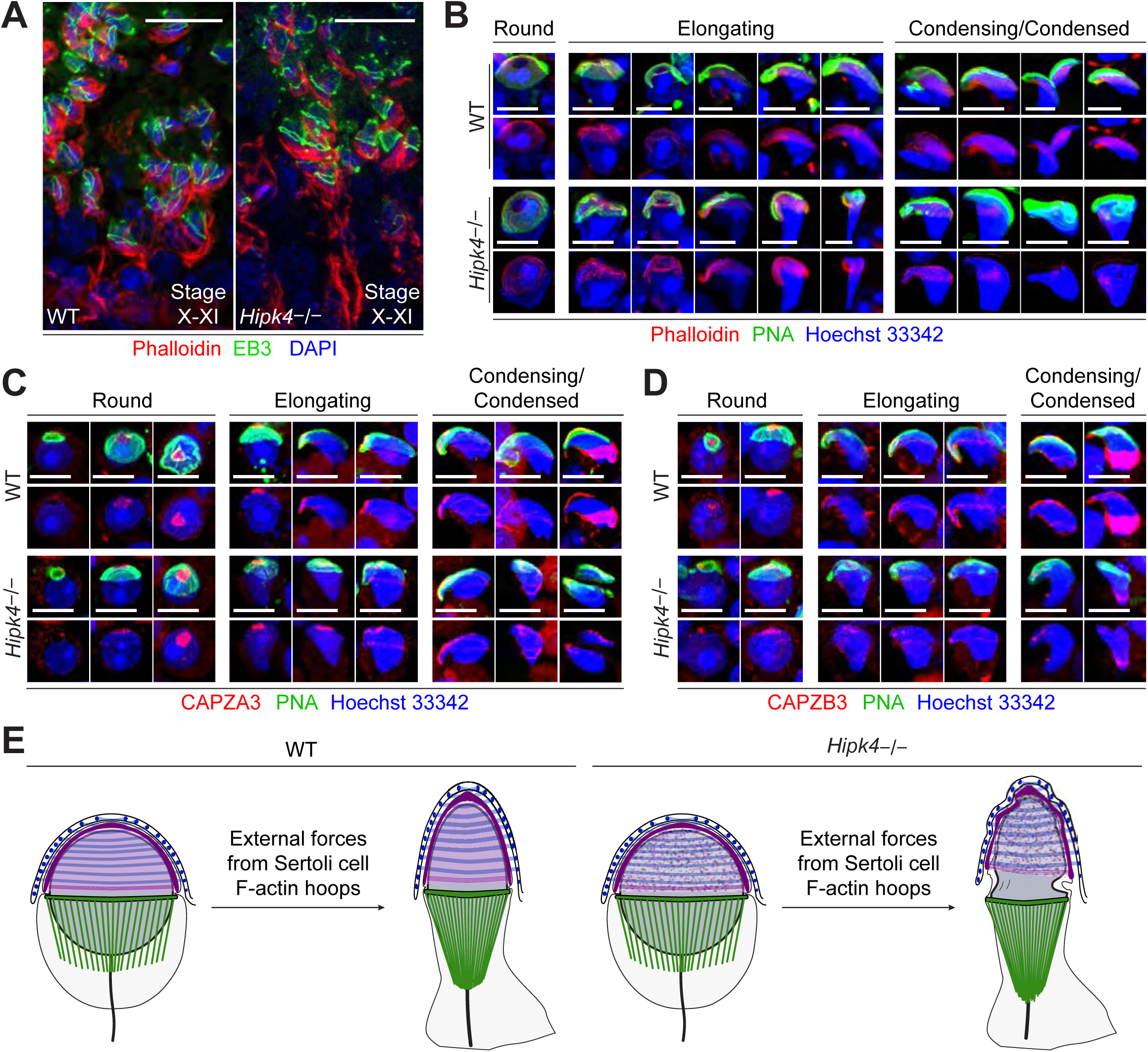
Loss of HIPK4 function alters F-actin dynamics in the acroplaxome. (A) Cryosectioned seminiferous tubules (stage X/XI) stained with Alexafluor-6 47-phalloidin and Alexafluor-48 8-anti-EB3 antibody to label basal and apical Sertoli cell ectoplasmic specializations. Nuclei were stained with DAPI. (B-D) Enzymatically dissociated elongating spermatids stained with FITC-PNA and Alexafluor-6 47-phalloidin (B), anti-CAPZA3 (C), or anti-CAPZB3 (D). Nuclei were stained with Hoeschst 33342. (E) A model for HIPK 4 function during spermiogenesis. Scale bars: A, 20 μm; B-D, 2 μm.

## DISCUSSION

HIPK4 is a dual-specificity kinase that is predominantly expressed in differentiating spermatids, and it has been previously reported that HIPK4 deficiency in mice can alter sperm morphology and number. Our studies reveal both the cell biological basis and reproductive consequences of these spermatogenic phenotypes. Our findings establish HIPK4 as an important regulator of the acrosome-acroplaxome complex during spermiogenesis. *Hipk4*^−/−^ spermatids form abnormal acroplaxome structures, and this F-actin- and keratin-containing plate becomes uncoupled from the Golgi-derived acrosome, most overtly at the groove belt in these mutant cells. HIPK4 function is also required to coordinate acrosome-acroplaxome and manchette dynamics during spermatid elongation, as evidenced by the grossly enlarged groove belt in elongating *Hipk4*^−/−^ spermatids. The resulting spermatozoa have misshapen heads, and a large fraction exhibit irregular tail morphologies. *Hipk4*^−/−^ males have lower concentrations of epididymal sperm, and these mutant germ cells have lower motility than their wild-type counterparts. Together, these spermatogenic defects closely mirror those observed in men with severe OAT syndrome, and like these clinical cases, HIPK4-deficient male mice are sterile.

Of these phenotypes, the abnormal head structures of HIPK4-deficient sperm are likely a dominant cause of sterility. Sperm produced by heterozygous *Hipk4* mutant males have lower than normal epididymal concentrations of sperm and reduced total motility, but the fertility of these animals is comparable to that of wild-type mice. Only homozygous *Hipk4* mutant germ cells exhibit head defects that become increasingly overt during spermiogenesis. Consistent with diminished head functions, *Hipk4*^−/−^ sperm are also incompetent for IVF and have reduced binding and penetration of COCs. This diminished fertility could reflect structural defects that physically abrogate sperm-egg interactions and/or concomitant perturbations that disrupt molecular processes within the head. The ability of *Hipk4*^−/−^ sperm to undergo capacitation and the acrosome reaction *in vitro* suggests that key intracellular biochemical pathways remain intact. Nevertheless, we cannot rule out the possibility that HIPK4 is required for these processes *in vivo*. The reduced motility of *Hipk4*^−/−^ sperm could also affect fertility within the female reproductive tract.

Our investigations reveal potential mechanisms by which HIPK4 could regulate the acrosome-acroplaxome complex. HIPK1-3 have been shown to directly phosphorylate homeodomain transcription factors (*Kim, Y. H., et al., 1998*; *Rinaldo, et al., 2008*), and the structurally related dual-specificity tyrosine-regulated kinase 1a (DYRK1A) targets various transcriptional activators and repressors (*Di Vona, et al., 2015*; *Litovchick, et al., 2011*; *Mao, et al., 2002*; *Woods, et al., 2001*; *Yang, et al., 2001*). Although it has been reported that HIPK4 can phosphorylate p53 *in vitro* (*Arai, et al., 2007*; *He, et al., 2010*), it is unlikely that HIPK4 acts primarily through transcriptional control. In contrast to other HIPK family members, HIPK4 lacks the homeobox-interacting domain and a nuclear localization sequence, and accordingly it localizes to the cytoplasm rather than the nucleus (*van der Laden, et al., 2015*). Moreover, our genome-wide microarray analyses reveal that loss of HIPK4 function does not lead to major transcriptional changes within the testis. These observations suggest that HIPK4 regulates the acrosome-acroplaxome complex through more direct biochemical mechanisms, and our cell biological studies strongly implicate HIPK4 in F-actin remodeling. HIPK4 expression in cultured somatic cells promotes the formation of branched F-actin structures and alters the phosphorylation state of multiple actin-crosslinking proteins. HIPK4 could similarly modulate F-actin networks within the acroplaxome. Consistent with this model, HIPK4 deficiency disrupts the levels of F-actin and actin-capping proteins localized to the acroplaxome marginal ring in condensing spermatids (CAPZA3 and CAPZB3), perhaps due to cytoskeletal dysregulation at earlier stages. We further note that transcripts encoding actin cytoskeleton components are among the few genes that are expressed in a HIPK4-dependent manner within the testis (Figure 5 – Source data 1). For example, the actin-related protein *Actr6* and the actin-capping protein *Capza2* were downregulated in *Hipk4*^−/−^ testis tissues, and upregulated genes includes those that encode actin-membrane crosslinkers (*Tln2*, *Flnb*, and *Ank2*), actin-based motors (*Myo5b* and *Myo10*), and components of the acrosome matrix (*Zp3r* and *Zan*). We speculate that these transcriptional changes reflect cellular responses to cytoskeletal defects caused by HIPK4 deficiency.

Based on our findings, we propose a model in which HIPK4 promotes F-actin remodeling in the acroplaxome, modulating its biophysical properties and interactions with the acrosome (Figure 7E). We hypothesize that loss of HIPK4 function alters acroplaxome stability and dynamics, rendering this cytoskeletal plate less able to distribute external forces applied by the Sertoli cell F-actin hoops. As a result, the spermatid nucleus fails to elongate properly, and the head structure becomes grossly misshapen. HIPK4 deficiency also uncouples the acrosome-acroplaxome complex at the groove belt, leading to severe anterior malformations that may disrupt the expulsion of excess cytoplasm and other aspects of spermatid differentiation. It is likely that germ cells with such significant defects are less competent for spermiation and less viable after their release into seminiferous tubule lumen, which could account for the lower concentrations of epididymal sperm in *Hipk4*^−/−^ mice.

HIPK4 joins the small collection of protein kinases that are known to be enriched or exclusively expressed in haploid male germ cells and required for spermatogenesis. For example, the kinases CSNK2A2, SSTK, and CAMKIV function in the spermatid nucleus and promote the histone-to-protamine transition (*Escalier, et al., 2003*; *Spiridonov, et al., 2005*; *Wu, et al., 2000*). TSSK1 and 2 are testis-specific kinases that regulate the chromatoid body, centrioles, and the developing sperm flagellum (*Jha, et al., 2013*; *Kueng, et al., 1997*; *Shang, P., et al., 2010*; *Xu, et al., 2008*), and TSSK4 and TSSK5 localize to the flagellum and control sperm motility (*Wang, et al., 2016*; *Wang, et al., 2015*). The only kinase known to be required for spermatid head shaping is TR-KIT, a splice variant of c-KIT that regulates manchette formation and later has a role in oocyte activation upon fertilization. A truncated, testis-specific version of FER, FERT, phosphorylates cortactin in the acroplaxome (*Keshet, et al., 1990*; *Kierszenbaum, et al., 2008*), but male mice lacking FERT are still fertile (*Craig, et al., 2001*). To the best of our knowledge, HIPK4 is the first kinase known to be essential for acrosome-acroplaxome function and male fertility.

Given the druggable nature of kinases, HIPK4 is also a new potential target for male contraception. Small-molecule HIPK4-specific antagonists could recapitulate the male sterility caused by disrupting the *Hipk4* gene in mice. Such treatments would reduce male fertility without interfering with the hypothalamus-pituitary-gonadal axis, damaging the testis or germline, or introducing genetic abnormalities to offspring if fertilization is achieved. Since HIPK4 expression is restricted to the final stages of sperm development, HIPK4 inhibitors also would induce sterility more quickly than drugs that perturb earlier steps in spermatogenesis. Contraceptive reversibility would be equally rapid. Moreover, HIPK4 antagonists could have minimal non-reproductive effects, as *Hipk4*^−/−^ mice appear to have otherwise normal physiology. Further investigations of HIPK4 therefore could not only elucidate the mechanisms that drive spermatid differentiation but also address a longstanding unmet need in reproductive medicine.

## MATERIALS AND METHODS

### Ethics statement

All animal studies were conducted in compliance with the Stanford University Institutional Animal Care and Use Committee under Protocol 29999. Vertebrate research at the Stanford University School of Medicine is supervised by the Department of Comparative Medicine’s Veterinary Service Center. Stanford University animal facilities meet federal, state, and local guidelines for laboratory animal care and are accredited by the Association for the Assessment and Accreditation of Laboratory Animal Care International. De-identified human testis sections were obtained from Stanford University in compliance with protocol IRB-32801.

### Animal use

*Hipk4^+/tm1b^* breeding pairs (C57BL/6NJ background, Stock No. 025579) and wild-type (C57BL/6NJ, Stock No. 005304) mice were purchased from The Jackson Laboratory. Mice used for this study were weaned at 19-22 dpp and genotyped using Platinum *Taq* DNA Polymerase (Invitrogen) and the following primers: wild-type forward 5’-CCTTTGGCCTTATACATGCAC-3’, wild-type reverse 5’-CAGGTGTCAGGTCTGGCTCT-3’, mutant forward 5’-CGGTCGCTACCATT ACCAGT-3’, mutant reverse 5’-ACCTTGAGATGACCCTCCTG-3’.

### Tissue distribution of Hipk4

TaqMan primers (Applied Biosystems) for *Hipk4* (Mm01156517_g1) were used to probe the Origene TissueScan^TM^ Mouse Normal cDNA Array according to the manufacturer’s protocols. Gene expression levels were normalized to *GAPDH* (Mm99999915_g1).

### In situ hybridization analysis

Whole testes were dissected and immediately fixed in freshly prepared modified Davidson’s fixative (30% formaldehyde,15% ethanol, 5% glacial acetic acid, 50% distilled water) for 16-20 hours and then washed and stored in 70% ethanol until further use. For *in situ* hybridization analyses, the fixed tissues were paraffin-embedded, cut into 10-μm sections, and mounted on slides. To detect *Hipk4* transcripts, we used the RNAscope^®^ 2.5 HD Detection Kit (Advanced Cell Diagnostics) with *Hipk4* probes (Mm-Hipk4, 428071), following the manufacturer’s protocol for formalin-fixed, paraffin-embedded (FFPE) sections. Hybridization probes for *Ppib* (BA-Mm-Ppib-1ZZ, 313911) and *dapB* (BA-DapB-1ZZ, 310043) were used as positive and negative controls, respectively.

### Assessment of fecundity

To test fertility by mating, we paired ≥ 7-week-old males (wild-type, *Hipk4*^+/–^, and *Hipk4*^−/−^) with age-matched, wild-type females for 18–21 days, and the number of live-born pups for each pairing was recorded. Male and female mice were paired in this manner for 2-4 rounds (6-12 weeks).

IVF and ICSI procedures were performed at the Transgenic, Knockout, Tumor Model Center at Stanford University. For IVF studies, C57BL/6NJ females were superovulated by injection with PMSG (ProSpec, HOR-272, 5U, 61-63 hours prior to oocyte-harvesting) and hCG (ProSpec, HOR-250, 5U, 48 hours after PMSG injection). On the day of the experiment, epididymal sperm were isolated using a “swim-out” method in TYH medium (120 mM NaCl, 5 mM KCl, 2.5 mM MgSO4, 1.0 mM KH2PO4, 25 mM NaHCO3, 2.5 mM CaCl2, 1 mM sodium pyruvate, 1.0 mg/mL glucose, 1.0 mg/mL methyl-β-cyclodextrin, and 1.0 mg/mL polyvinylalcohol; pH 7.2) and transferred to TYH medium containing bovine serum albumin (BSA; final concentration of 5 µg/mL). Cumulus-oocyte complexes (COCs) were then harvested from the oviducts of these superovulated females in M2 medium with HEPES buffer (Sigma, M7167), and transferred through three 50 µL drops of HTF medium (Millipore, MR-070-D) containing 0.25 mM reduced glutathione under mineral oil, pre-equilibrated to 37 °C, 5% CO2. After 1 hour of capacitation, motile sperm (∼ 5.0 × 10^5^) were added to the drop of HTF medium containing COCs and incubated for 4 hours at 37 °C and 5% CO2. Sperm-oocyte complexes were then washed four times in M2 medium and incubated in 30 µL KSOM medium (Millipore, MR-101-D) at 37 °C, 5% CO_2_ under mineral oil. The number of two-cell, morula, and blastocyst-stage embryos were then counted over the next 72 hours.

ICSI experiments were performed as previously described (*Yoshida, et al., 2007*). Briefly, motile sperm were harvested from the epididymis, and sperm heads were injected into the cytoplasm of CD1 oocytes using a piezo-actuated micromanipulator. The injected embryos were cultured in KSOM medium at 37 °C. After 24 hours, live two-cell embryos were either cultured until the blastocyst stage or implanted into the oviducts of pseudo-pregnant female mice.

### Antibodies

The following primary antibodies were used for western blotting and immunofluorescence imaging: anti-HIPK4 (FabGennix International, rabbit pAb generated against the peptide sequence PAGSKSDSNFSNLIRLSQVSPED, lot 1651.Pb1.AP); anti-KPNB1 (H-300) (Santa Cruz Biotechnology, sc-11367, rabbit pAb); anti-phosphotyrosine (Upstate/Millipore, 4G10^®^ Platinum, 05-1050, lot 2723728, rabbit pAb); anti-ZP3R/mouse sp56 (7C5) (QED Bioscience, 55101, lot 051614-120816, mouse mAb); anti-IZUMO1 (125) (Abcam, ab211626, lot GR279965-4, rat mAb); anti-SPACA1 (Abcam, ab191843, lot GR312512-3, rabbit pAb); anti-FLAG^®^ (M2, Sigma, F3165, mouse mAb); anti-alpha tubulin (3H3085) (Santa Cruz Biotechnology, sc-69970, lot G0109, rat mAb); anti-CAPZA3 (Progen, GP-SH4, lot 804091, guinea pig pAb); anti-CAPZB3 (Progen, GP-SH5, lot 804081, guinea pig pAb); anti-DPY19L2 (gift from Christophe Arnoult, rabbit pAb), anti-SUN1 (gift from Manfred Alsheimer, guinea pig pAb); anti-nesprin3 (gift from Arnoud Sonnenberg, rabbit pAb); Alexa Fluor^TM^ 488-conjugated anti-EB3 (EPR11421-B) (Abcam, ab203264, lot GR227133-1, rabbit pAb).

Secondary antibodies included HRP-conjugated sheep anti-mouse IgG (GE Healthcare, NA931V, lot 9682503), HRP-conjugated donkey anti-rabbit IgG (GE Healthcare, NA934V, lot 9780721), Alexa Fluor^TM^ 594-conjugated anti-guinea pig (Invitrogen, A11076, lot 1924784), Alexa Fluor^TM^ Plus 647-conjugated goat anti-mouse IgG (Invitrogen, A32728, lot UB275580), Alexa Fluor^TM^ 647-conjugated goat anti-rat IgG (Invitrogen, A21247, lot 37177A), and Alexa Fluor^TM^ Plus 647-conjugated goat anti-rabbit IgG (Invitrogen, A32733, lot TL272452).

### Western blot analysis

To detect HIPK4 protein, testis lysates were prepared by sonication of ∼100-mg pieces of freshly removed whole testis in 1.0 mL ice-cold RIPA buffer containing protease inhibitors (cOmplete^TM^, EDTA-free Protease Inhibitor Cocktail Tablets, Roche, 4693159001) and phosphatase inhibitors (PhosSTOP^TM^, Roche, 4906837001). Equivalent amounts (12 µg) of total protein per sample were diluted with 6x Laemmli sample buffer, boiled for 5 minutes, and loaded on a 3-8% tris-acetate gel (Bio-Rad) for SDS-PAGE. Proteins were transferred to a PVDF membrane and immunoblotted with the following primary antibodies: anti-HIPK4 [0.55 μg/mL, 1:1000 dilution in phosphate buffered saline (PBS) containing 0.1% Tween 20 and 4% BSA] or anti-KPNB1 (1:1000). Chemiluminescence detection was conducted with HRP-conjugated secondary antibodies and either SuperSignal^TM^ West Dura or SuperSignal^TM^ Femto kits (Pierce/Thermo Fisher Scientific, 34076 or 34095, respectively).

### Assessment of sperm quality

Epididymides from male mice (8-9 weeks old) were dissected and cleared of fat tissue before being cut open with fine scissors in 200 µL HTF medium equilibrated at 37 °C and 5% CO2. The sperm suspension (10 µL) was counted and assessed for total motility at 100x magnification using Leja slides and a microscope equipped with a reticle. Sperm quality parameters of males (15-17 weeks old) were also assessed by the Mouse Biology Program at UC-Davis using the following procedures. Morphology was assessed visually, and CASA was used to confirm sperm concentrations and motility. Sperm were viewed randomly under 400x or 1000x magnification to determine the percentage with abnormal head sizes and shapes (macrocephalous, microcephalous, tapered, triangular, olive, pin, banana, amorphous, collapsed, abnormal hook, irregularly shaped, etc.) or abnormal midpieces or tails (bent, coiled, short, thin, crinkles, irregularly shaped, etc.). Sperm were not included in the morphology assessment if they were: (1) aggregated; (2) had a back or abdomen view of the head; or (3) were decapitated. At least 100 sperm in 5 or more fields of view were evaluated for each experimental condition.

### Assessment of sperm maturation and function

To assess capacitation by western blot, motile sperm were collected in 2.5 mL of TYH medium (BSA-free, pH 7.4, and CO2-equilibrated) from cauda epididymides using the “swim out” method at 37 °C and 5% CO_2_. The sperm suspension was then centrifuged at 600 *g*, and all but 0.4 mL of the supernatant was removed to achieve a final concentration of ∼40 × 10^6^ cells/mL. In fresh polystyrene tubes, 200 µL of the sperm solution was added to 1.0 mL TYH medium containing BSA (10 mg/mL), and the suspension was incubated at 37 °C for 1.5 hours. The sperm were then centrifuged at 13,000 *g* for 1 minute, washed with PBS containing phosphatase inhibitors, and pelleted once more. After the addition of 1x Laemmli sample buffer without reducing agent (25 µL), and the samples were boiled for 5 minutes and centrifuged at 13,000 *g*. The supernatants were collected and 2 µL of β-mercaptoethanol was added to each sample, which were then boiled again for 1 minute. Following SDS-PAGE, proteins were transferred to a PVDF membrane and immunoblotted with an anti-phosphotyrosine antibody (1:1000 dilution, Upstate/Millipore) for chemiluminescence detection.

To perform *in vitro* acrosome reactions, sperm were collected and capacitated as described above. Following 1.5 hours of capacitation, the Ca^2+^ ionophore A23187 (hemicalcium salt; Sigma, |C9275; 1000x stock dissolved in ethanol) was added to the suspensions to achieve final concentrations of 10 µM. After incubation for 1 hour at 37 °C and 5% CO2, and the cells were pelleted by centrifugation at 600 *g*, and all but 0.5 mL of the supernatant was removed. The following steps were then performed in microcentrifuge tubes, and the cells were pelleted at 600 *g* and washed with PBS between each step. Sperm were fixed by adding 2.0 mL 4% paraformaldehyde in PBS for 15 minutes and permeabilizing with 0.3% Triton-X100 in PBS for 5 minutes. For experiments measuring acrosome exocytosis, the sperm were then incubated for 30 minutes in PBS containing 2% BSA, 0.01% Triton X-100, and fluorescein-conjugated peanut agglutinin (FITC-PNA, 10 µg/mL, Sigma, L7381, lot 046M4030V) and 10 µg/mL Hoechst 33342 nuclear stain. After washing the cells with PBS, they were mounted on microslides with Vectashield Vibrance medium (Vector Labs, H-1700, ZE1011). For immunolabeling experiments, the cells were then divided into separate tubes, blocked for 1 hour with PBS containing 2% BSA and 0.01% Triton X-100, incubated for 1 hour at room temperature with primary antibodies (10 μg/mL final concentration in blocking buffer), and then washed once with blocking buffer. The samples were then incubated with fluorescently labeled secondary antibodies and 10 µg/mL FITC-PNA in blocking buffer for 30 minutes. The sperm were washed once more, transferred to microscope slides, mounted with Prolong Gold medium with DAPI, and then imaged on a Zeiss LSM 700 confocal microscope equipped with a 63x oil-immersion objective. ImageJ software (NIH) was used to create maximum-intensity Z-stack projections, and Photoshop CS6 (Adobe) was used to crop images and adjust fluorescence intensity levels.

To assay for oocyte binding *in vitro*, we followed the same general protocol described for IVF experiments. After 3 hours of sperm-COC incubation and five washing steps, we transferred the complexes in a small volume into a 50-µL drop of PBS containing 4% paraformaldehyde on a microscope slide. After 30 minutes, excess liquid was carefully removed with a Kimwipe, and the slides were mounted with Prolong Gold medium with DAPI. Fluorescence and DIC imaging on a Leica DMi8 microscope at 200x magnification were used to assess the number of sperm bound to oocytes.

### TUNEL assays

To measure DNA double strand breaks by terminal deoxynucleotidyl transferase-mediated dUTP nick end labeling (TUNEL), we used the *In Situ* Cell Death Detection Kit (Sigma-Aldrich, 11684795910), following the manufacturer’s protocol (ver. 17) for cell suspensions. Fluorescence imaging was performed using a Zeiss LSM 700 confocal microscope equipped with a 63x oil-immersion objective. ImageJ software (NIH) was used to create maximum-intensity Z-stack projections and TUNEL-positive cells were manually counted (n = >357).

### Histology of testis and epididymis

Histological staining (PAS and H&E) was performed on formalin-fixed, paraffin-embedded testis and epididymis sections (10 µm). To obtain these sections, the tissues were removed from 12- to 15-week-old males, immediately fixed in modified Davidson’s fixative for 16 hours, and stored in 70% ethanol until paraffin-embedding, sectioning, and mounting on microscope slides. The slides then were heated for 1 hour at 50 °C, washed with xylenes, re-hydrated, and stained with periodic acid solution (Sigma, 3951) and Schiff’s reagent (Sigma, 3952016) and/or counterstained with modified Harris hematoxylin solution (Sigma, HHS32) and Eosin Y (Sigma, E4009).

### Electron microscopy

For scanning electron microscopy, epididymal sperm were collected in TYH medium by the “swim-out” method at 37 °C, pelleted at 200 *g*, resuspended in 0.1 M sodium cacodylate buffer (pH 7.4) containing 2% paraformaldehyde and 4% glutaraldehyde, and transferred to individual wells of a 24-well plate containing poly-D-lysine-coated 12-mm coverslips. The samples were allowed to fix overnight at 4 °C and then post-stained with 1% aqueous osmium tetroxide (EMS, 19100) for 1 hour. OsO4-treated samples were rinsed in ultrafiltered water three times and gradually dehydrated in increasing concentrations of ethanol (50%, 70%, 90%, 2 × 100%; 15 minutes each). Each coverslip was then dried at the critical point with liquid CO2 using a Tousimis Autosamdri®-815A system and 15-minute purge time. Dried samples were sputter-coated (100 Å, Au/Pd) before imaging with a Zeiss Sigma FE-SEM using In-Lens and lateral Secondary Electron detection at 3.02 kV.

For transmission electron microscopy, samples were fixed in 0.1 M sodium cacodylate buffer (pH 7.4) containing 2% glutaraldehyde and 4% paraformaldehyde at room temperature for 1 hour. The fixative was then replaced with cold aqueous 1% OsO4, and the samples were allowed to warm to room temperature for 2 hours, washed three times with ultrafiltered water, and stained in 1% uranyl acetate for 2 hours. Samples were then dehydrated in a series of ethanol washes, beginning at 50%, then 70% ethanol, and then moved to 4 °C overnight. They were then placed in cold 95% ethanol and allowed to warm to room temperature, changed to 100% ethanol for 15 minutes, and finally to propylene oxide (PO) for 15 minutes. Samples were next incubated with sequential EMbed-812 resin (EMS, 14120):PO mixtures of 1:2, 1:1, and 2:1 for 2 hours each and stored overnight in 2:1 resin:PO. The samples were then placed into 100% EMbed-812 resin for 4 hours, moved into molds with fresh resin, orientated and warmed to 65 °C overnight. Sections were taken between 75 and 90 nm, picked up on formvar/carbon-coated slot Cu grids, stained for 40 seconds in 3.5% uranyl acetate in 50% acetone, followed by staining in 0.2% lead citrate for six minutes. The sections were then imaged using a JEOL JEM-1400 120 kV instrument and a Gatan Orius 832 4k × 2.6k digital camera with 9-µm pixel size.

### Microarray assay

Three testes from 12-week old wild-type and *Hipk4*^−/−^ mice were snap frozen in liquid N2, and stored at −80 °C. Samples were thawed and 50-mg portions were immediately homogenized in TRIzol, and RNA was isolated using the Direct-zol^TM^ RNA MiniPrep Plus kit (Zymo Research, R2070S) and stored at −80 °C. RNA profiling was then conducted in quadruplicate Mouse Clariom^TM^ D assays (Applied Biosystems, 902514), following the manufacturer’s protocol. Briefly, 50 ng of each sample was used as an input into the GeneChip^TM^ WT Plus Reagent Kit (P/N 902281), and labeled targets were hybridized to arrays in a GeneChip™ Hybridization Oven 645 (00-0331). Washing and staining steps were performed on a GeneChip™ Fluidics Station 450 (P/N 00-0079), and arrays scanned on a GeneChip™ Scanner 3000 7G (P/N 00-210) system. Data were analyzed using the Transcriptome Analysis Console 4.0 software package.

### Retrovirus production

Murine *Hipk4* and the *K40S* mutant genes were obtained by PCR using the primers 5’-CAAAAAAGCAGGCTCAGCCACCATGGCCACCATCCAGTCAGAGACTG-3’ and 5’-CAAGAAA GCTGGGTCGTGGTGCCCTCCAACATGCTGCAG-3’ for wild-type *Hipk4* and 5’-TCGATCCTGA AGAACGATGCGTACCGAAGC-3’ and 5’-GATGGCCACCATTTCACCTGTACTCCGAC-3’ for the *Hipk4-K40S* mutant. pCL-ECO was purchased from Imgenex, and pBMN-I-GFP was provided by Gary Nolan. For Gateway recombination-mediated cloning, the PCR products were amplified further with the primers 5’-GGGGACAAGTTTGTACAAAAAAGCAGGCTCA-3’ and 5’-GGGGACC ACTTTGTACAAGAAAGCTGGGTC-3’ and Phusion polymerase (New England Biolabs) to add attB adapter sequences. The clones were then transferred into pDONR223 in a BP recombination reaction using BP clonase II (Invitrogen) according to the manufacturer’s protocols. pDONR223 entry constructs were next transferred to pBMN-3xFLAG-IRES-mCherry-DEST vectors using LR clonase II (Invitrogen) according to the manufacturer’s protocols. pBMN-HIPK4-Y175F-3xFLAG-IRES-mCherry was later generated by site-directed mutagenesis of the wild type construct using the primers 5’-CGCTATGTGAAGGAGCCTTTCATCCAGTCCCGCTTC TAC-3’ and 5’-GTAGAA GCGGGACTGGATGAAAGGCTCCTTCACATAGCG-3’ and *Pfu*Ultra II Fusion polymerase (Agilent).

Retroviral stocks were prepared from HEK-293T cells seeded in 10-cm tissue culture dishes (∼4 × 10^6^ cells/dish) in 10 mL of culture medium. Approximately 18 hours post-seeding, each 10-cm dish was transfected as follows: pBMN-3xFLAG-IRES-mCherry-DEST plasmids containing wild type or mutant *Hipk4* (7.4 µg) and the pCL-ECO retrovirus packing vector (4.4 µg) were diluted in OMEM medium (375 µL). This DNA mixture was added to 40 µL Fugene® HD reagent (Promega) in OMEM (335 µL) and incubated at room temperature for 15 minutes, before being gently added to culture medium on cells. After 24 hours, the medium was replaced with DMEM containing 10 mM HEPES (pH 7.4), 3% fetal bovine serum, 7% calf serum, and 1% sodium pyruvate. Retrovirus-containing supernatant was then collected three times at 24-hour intervals, passed through a 0.45-µm filter, and stored at −80 °C.

### Quantitative phosphoproteomics by mass spectrometry

To obtain peptides suitable for phospho-enrichment and mass spectrometry studies, NIH-3T3 cells (passage 6) were seeded in 10-cm dishes (five per condition) at 1.5 × 10^6^ cells/dish with DMEM containing 10% calf serum, 0.1% sodium pyruvate, 100 U/mL penicillin, and 0.1 mg/mL streptomycin and transduced after 12 hours with the appropriate retrovirus and polybrene (8 µg/mL) to achieve a multiplicity of infection (MOI) >4. After 48 hours, the culture medium was replaced with DMEM containing 0.1% CS for 4 hours, and then all dishes were transferred to a cold room, rinsed twice with ice-cold PBS, and incubated on a rocker for 10 minutes with 0.4 mL RIPA lysis buffer containing protease inhibitors (cOmplete^TM^ tablets, Roche) and phosphatase inhibitors (PhosSTOP^TM^ tablets, Roche; 0.2 mM PMSF, Sigma). Cells were manually scraped off of each dish, and the suspensions were transferred to 15-mL Falcon tubes on ice. Cells were sonicated for 30 seconds on ice, and the lysates were cleared by centrifugation at 14000 *g*. A portion of each lysate was removed for protein concentration determination and western blot analyses. The remaining protein lysates were precipitated with 14 mL of cold acetone at −80 °C. Precipitates were pelleted, the supernatant was thoroughly removed, and proteins were resolubilized with 2.0 mL of 8.0 M urea, 50 mM sodium bicarbonate (pH 8.0). Samples were reduced at room temperature for 30 minutes with the addition of DTT to a final concentration of 5 mM, and then alkylated in the dark using acrylamide at a final concentration of 10 mM for 30 minutes. To digest proteins, samples were diluted to 1 M urea with 50 mM sodium bicarbonate (pH 8.0), Protease Max Surfactant (Promega, V2072) was added to a final concentration of 0.03%, and a Trypsin–LysC protease mix (Promega, V5073) was added at ∼1:40 ratio to the protein concentration of each sample. Samples were incubated at 37 °C for 14 hours.

The digests were acidified to pH 3.5 with formic acid and incubated for 15 minutes at 37 °C to break down surfactants. Peptides were purified using Oasis HLB columns (3 mL) (Waters Co., WAT094226) according to the manufacturer’s protocol, lyophilized overnight, and resolubilized in a 4:1 acetonitrile/H_2_O mixture containing 0.1% trifluoracetic acid. Peptide concentrations were determined using the Pierce Peptide Quantification Colorimetric Assay. Samples were adjusted to 1.0 mg/mL by adding a 4:1 acetonitrile/H2O mixture containing 0.2% trifluoracetic acid, and 20 µL of the resulting solution was saved for “total peptide” mass spectrometry runs. A standard set of phosphopeptides (4 pmol/peptide sample, MS PhosphoMix1 Light, Sigma, MSPL1) were spiked into each sample. Phosphopeptide-enrichment steps were performed using a 1:1 mix of MagReSyn® Ti-IMAC and MagReSyn® TiO2 resins (ReSyn, MR-TIM005 and MR-TID005) according to the manufacturer’s protocols. The eluted peptides were acidified to pH 2.5 with 10% TFA in H2O and reduced to a volume of 5-10 µL using a SpeedVac concentrator. The samples were further enriched for hydrophilic peptides by purification through graphite spin columns (Pierce, 88302) according to the manufacturer’s protocol. Samples were adjusted to pH ∼8.0 with a 100 mM triethylammonium bicarbonate solution and dried on a SpeedVac concentrator. The peptides from different conditions were then isobarically labeled using the TMT-6plex kit (Pierce, 90061) according to the manufacturer’s protocols and pooled together. Finally, these pooled peptide samples were fractionated into six fractions using a high-pH reversed-phase peptide fractionation kit (Pierce, 84868, lot RF231823B), and each fraction was run on a Thermo Orbitrap Fusion Tribrid for LC-MS/MS analysis.

Peptides were identified using SEQUEST software, and individual species were removed from the analysis if they: (1) had a false discovery rate was above 1%; (2) were contaminating peptides (i.e. bovine or human); (3) were not phosphorylated. The signal intensities were then normalized for each TMT channel, based on the sum of the signals detected for the standard phosphopeptide mix that had been spiked into each sample prior to the phospho-enrichment step. Peptides that were identified as having the same phosphorylation state were combined, converging on 6,947 phosphosites with their associated A scores. The fraction of individual TMT signals relative to the total intensity for each peptide was then determined.

### Immunofluorescence studies

To determine the effects of *Hipk4* overexpression in NIH-3T3 cells, the fibroblasts were seeded into 6-well plates at a density of 150,000 cells/well and infected 18 hours later with the appropriate retrovirus at an MOI of ∼4. One day after infection, cells were reseeded into 24-well plates containing poly-D-lysine-coated 12-mm glass coverslips and cultured for 24 hours in DMEM containing 10% calf serum, 100 U/mL penicillin, and 0.1 mg/mL streptomycin. The cells were then treated with DMEM containing 0.5% calf serum and the antibiotics for an additional 24 hours, fixed in PBS containing 4% paraformaldehyde for 10 minutes at room temperature, and washed 3 × 5 minutes with PBS. The cells were permeabilized with PBS containing 0.3% Triton X-100 for 5 minutes and blocked for 2 hours at 4 °C in PBS containing 2% BSA and 0.1% Triton X-100. The cells were then incubated for 1 hour at room temperature with primary antibodies (1:200 dilution in blocking buffer), washed 3 × 5 minutes with PBS, incubated for 30 minutes with the appropriate secondary antibodies (1:400 dilution in blocking buffer) and/or Alexa Fluor^TM^-647 conjugated phalloidin (1:400, Invitrogen, A22287, lot 1884190), and washed twice more with PBS. Nuclei were stained with DAPI, and the coverslips were rinsed briefly in water and mounted onto slides using Prolong Gold Antifade reagent (Invitrogen).

Immunofluorescence imaging of testis sections was conducted using cryosections of fresh frozen tissue (10 μm thick). Once sectioned, samples were fixed on the slides with PBS containing 4% paraformaldehyde for 30 minutes at room temperature, permeabilized with PBS containing 1% Triton-X-100 for 15 minutes, blocked with PBS containing 2% BSA and 0.01% Triton X-100, incubated with primary antibody (10 μg/mL in blocking buffer) for 2 hours, washed three times with PBS containing 0.01% Triton X-100, incubated with secondary antibody (1:400 dilution in blocking buffer) along with either Alexa Fluor^TM^-conjugated phalloidin (1:400) or fluorescently labeled anti-EB3 (1:100), for 30-45 minutes, washed three times with PBS containing 0.01% Triton X-100, and mounted with coverslips with Prolong Gold with DAPI.

Immunofluorescence imaging of isolated spermatids was conducted on cells enzymatically dissociated from seminiferous tubules. Briefly, testes were dissected and decapsulated in warm DMEM containing HEPES (10 mM, pH 7.4). Collagenase Type I (final concentration of 1.5 mg/mL, Worthington Biochemical, LS004194, lot SF8B18091A) and DNase I (1.0 mg/mL) were added, and tubules were gently separated and incubated at 37 °C for 15 minutes. The tubules were then transferred to DMEM, cut into small fragments using fine scissors, and incubated with trypsin (2.0 mg/mL, Worthington Biochemical, LS003702) and DNase I (2.0 mg/mL) in DMEM for 20 minutes at 37 °C with vigorous physical mixing every 4 minutes using a plastic transfer pipette. BSA was added to stop enzymatic digestion, and the cells were pelleted at 400 *g* for 10 minutes at 4 °C. The cells were thoroughly resuspended in PBS containing 1 mg/mL polyvinyl alcohol (PVA) and carefully loaded onto a gradient column of BSA in DMEM containing HEPES buffer (10 mM, pH 7.4). The columns contained 25-mL zones of 4%, 3%, 2%, and 1% BSA. After 4 hours of gravity sedimentation, fractions of cells were collected, and those containing round, elongating, and condensed spermatids were combined and pelleted at 400 *g*. Cells were suspended in the PBS/PVA buffer, pelleted, suspended in a hypotonic sucrose solution (20 mM HEPES, 50 mM sucrose, 17 mM sodium citrate) for 10 minutes, pelleted, and then fixed in PBS containing 4% paraformaldehyde at room temperature for 15 minutes. The cells were then permeabilized with PBS containing 1 mg/mL PVA and 1.0% Triton X-100, blocked with PBS containing 2% BSA and 1 mg/mL PVA, and incubated with the appropriate primary antibody (1:50 dilution in blocking buffer) at 4 °C overnight with end-over-end rotation. After three 10-minute washes with PBS containing 0.01% Triton X-100 and 1 mg/mL PVA at room temperature, the cells were incubated with the appropriate fluorescently labeled secondary antibody (1:400 dilution in blocking buffer) along with FITC-PNA (1:1000), Alexa Fluor^TM^-647 conjugated phalloidin (1:400), and/or fluorescently labeled anti-EB3 (1:200) for colocalization) for 30-45 minutes, washed once, incubated with PBS/PVA buffer containing Hoechst 33342 dye for 10 minutes at room temperature, washed once more, and mounted on microscope slides using Vectashield Vibrance medium.

All fluorescence imaging was performed using a Zeiss LSM 700 or 800 confocal microscope equipped with a 63x oil-immersion objective. ImageJ software (NIH) was used to create maximum-intensity Z-stack projections, and Photoshop CS6 (Adobe) was used to crop images and adjust fluorescence intensity levels.

## SUPPLEMENTAL DATA

Figure 2 – Figure supplement 1

Figure 2 – Figure supplement 2

Figure 3 – Figure supplement 1

Figure 3 – Figure supplement 2

Figure 4 – Figure supplement 1

Figure 4 – Figure supplement 2

Figure 4 – Figure supplement 3

Figure 5 – Source data 1

Figure 6 – Source data 1

## ACKNOWLEDGMENTS

This work was supported by R21 HD78385 (J.K.C.), a Male Contraceptive Initiative Research Grant (J.K.C.), and postdoctoral fellowships from the American Cancer Society (J.A.C.) and the Male Contraceptive Initiative (J.A.C.). We gratefully acknowledge Chris Adams and Ryan Lieb (Stanford University Mass Spectrometry Core) for critical help with mass spectrometry experiments, and Lydia-Marie Joubert (Cell Sciences Imaging Facility, Stanford University) for providing training related to our scanning electron microscopy experiments. We are also indebted to Pablo Visconti (University of Massachusetts), Moira O’Bryan (Monash University), George Gerton (University of Pennsylvania), and Michael Eisenberg (Stanford University) for their thoughtful discussions and protocols. Antibody reagents were kindly provided by Christophe Arnoult (Université Grenoble Alpes), Manfred Alsheimer (University of Würzburg), and Arnoud Sonnenberg (Universiteit Leiden). John Higgins (Stanford University) supplied human testis sections. The electron microscopy studies were supported, in part, by ARRA Award Number 1S10RR026780-01 from the National Center for Research Resources (NCRR).

## AUTHOR CONTRIBUTIONS

J.A.C. designed and performed the experiments, analyzed the data, and wrote the paper. P.G.R. performed the qPCR and microarray assays. Z.J.H. performed animal husbandry, *in situ* hybridization, immunofluorescence, and SEM experiments. J.E.E. analyzed mass spectrometry data, J.J.P. prepared samples for TEM imaging, B.B. performed sperm concentration and percent motility assays. Y.L, J.L., and H.F. performed IVF and ICSI experiments and assisted with sperm-oocyte binding assays. J.K.C. designed the experiments, analyzed the data, and wrote the paper.

---

**Figure 2 – figure supplement 1.**
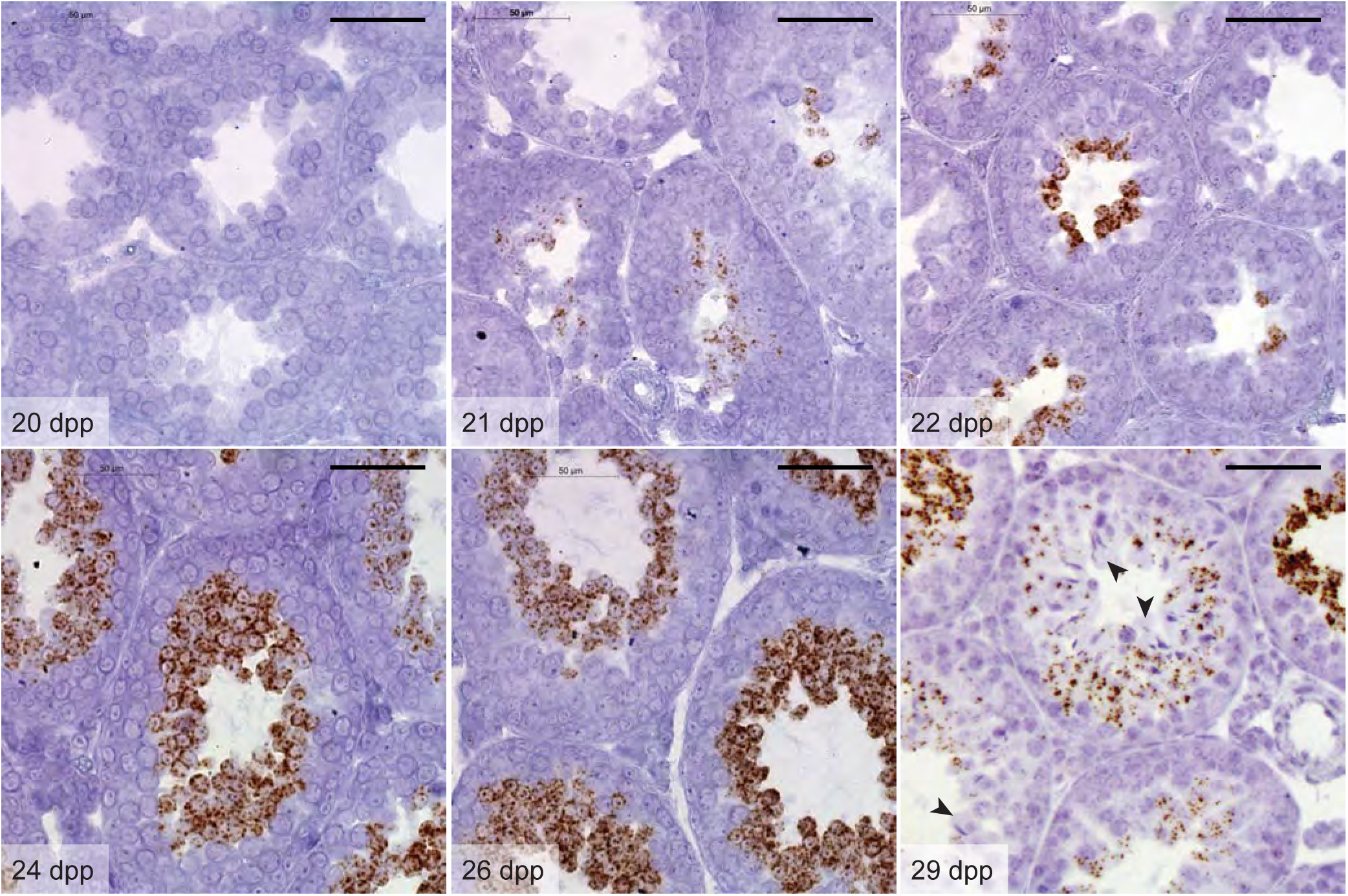
*Hipk4* mRNA expression during the first wave of murine spermatogenesis. *Hipk4* transcript labeling by *in situ* hybridization in formalin-fixed, paraffin-embedded testis sections from wild-type C57BL/6NJ mice of the designated ages. Arrowheads indicate the loss of *Hipk4* expression in elongating spermatids circumscribing the seminiferous tubule lumen (29 dpp). Scale bars: 50 µm.

**Figure 2 – figure supplement 2.**
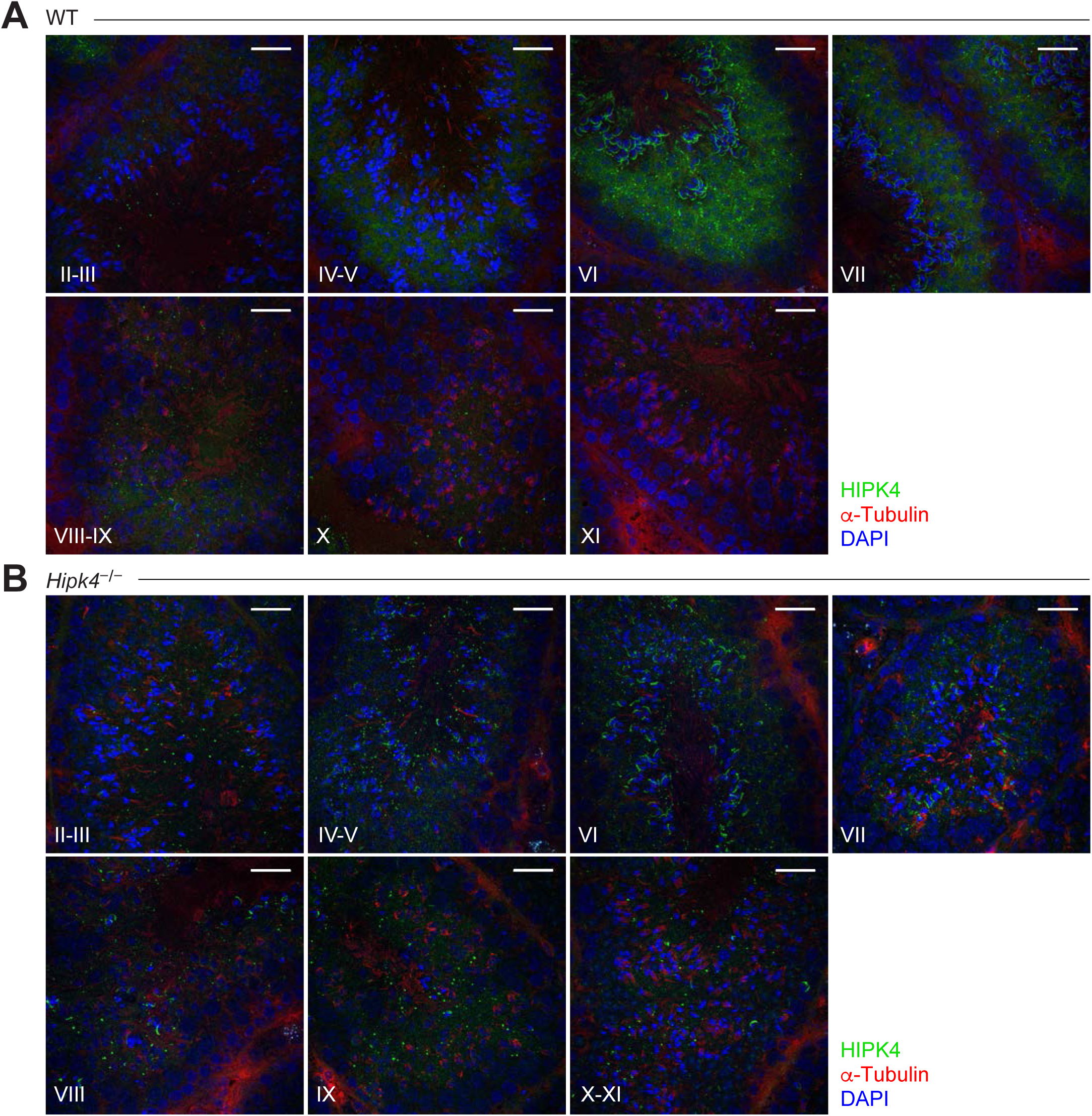
HIPK4 protein expression in adult murine seminiferous tubules. (A-B) Immuno-fluorescence images of cryosectioned testes from adult wild-type (A) and *Hipk4* knockout (B) mice. Stages for each testis section are shown. Scale bars: 20 µm.

**Figure 3 – figure supplement 1.**
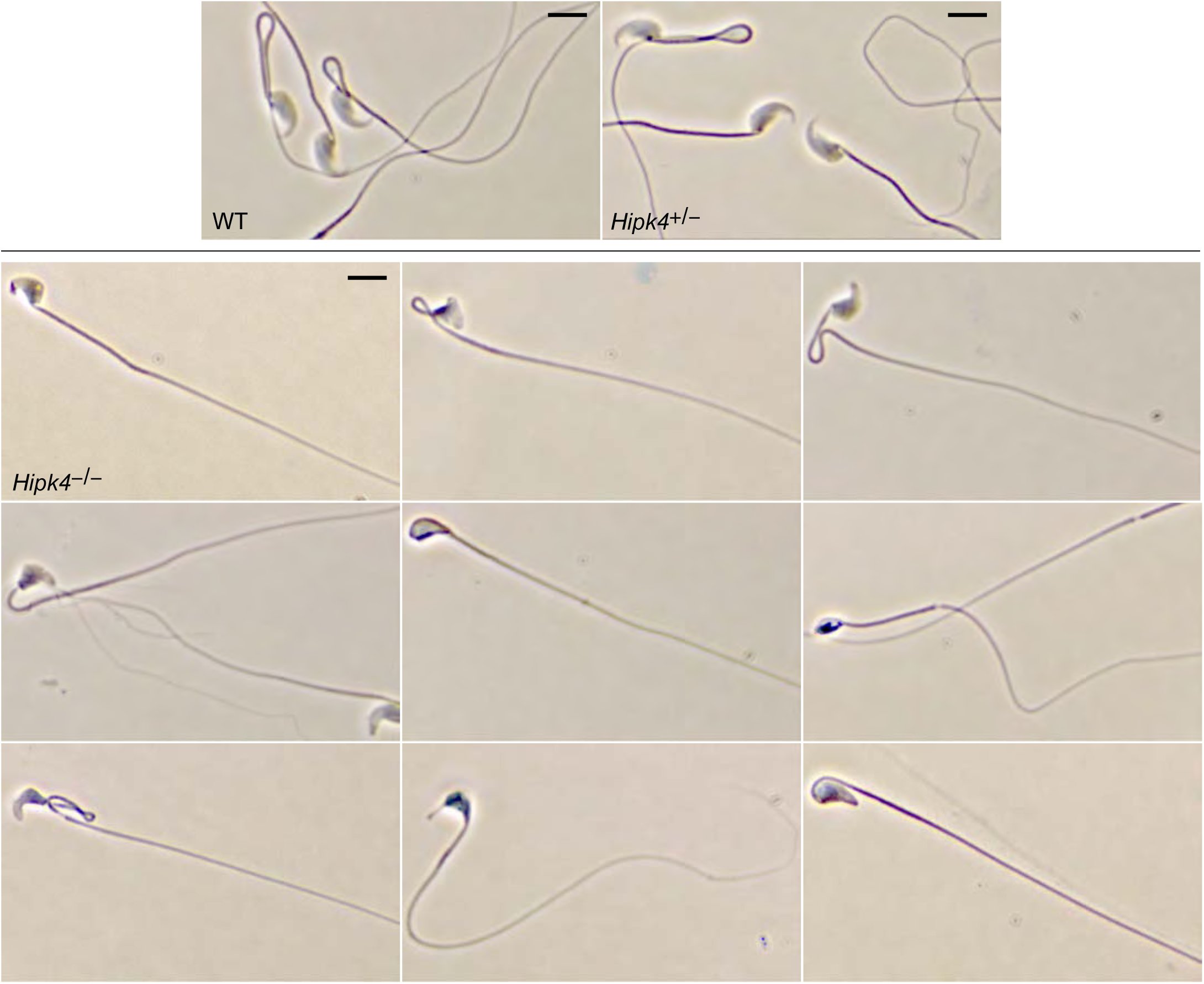
Morphologies of wild-type and *Hipk4* mutant sperm. Phase contrast images of sperm obtained from mice with the designated genotypes. Sperm were treated with Diff-Quik stain prior to imaging, and multiple examples of homozygous mutant sperm are shown to illustrate the range of morphological phenotypes. Scale bars: 2 µm.

**Figure 3 – figure supplement 2.**
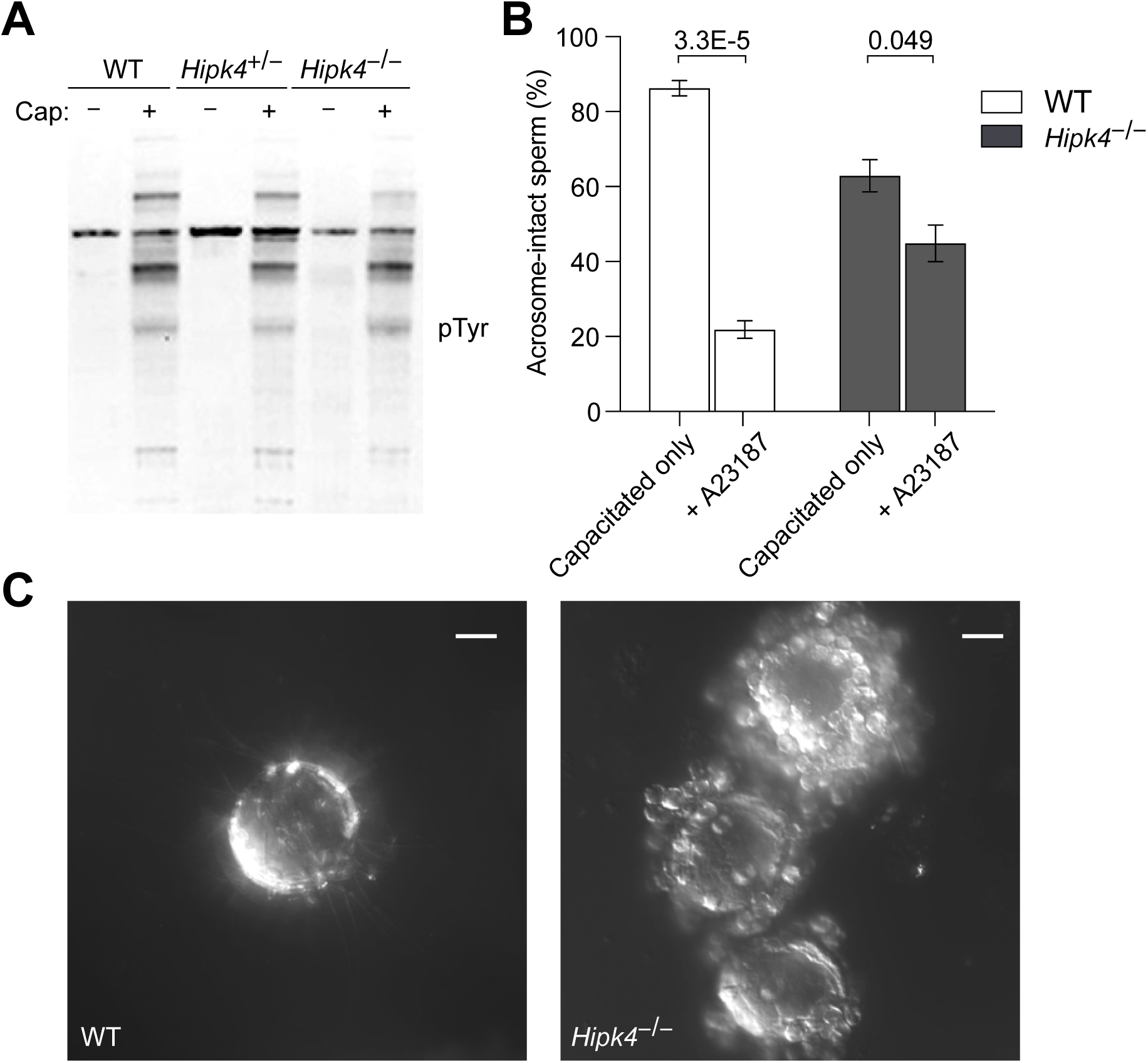
HIPK4 is not essential for sperm capacitation or acrosomal exocytosis. (A) Western blot detection of soluble phosphotyrosine-containing proteins in sperm before or after capacitation with TYH medium + BSA (Cap buffer). (B) Percentage of wild-type or *Hipk4* knock-out sperm with an acrosome after treatment with Cap buffer for 1.5 hours or after an additional 1-hour incubation with either the Ca^2+^ ionophore A23187. Data are the average of three independent experiments ± s.e.m., and the number of sperm counted for each condition was between 213-1228. *P*-values (T-test; two-tailed, equal variance) are shown. (C) Representative differential interference contrast (DIC) images of oocytes following standard IVF conditions using wild-type or *Hipk4* knockout sperm. Scale bars = 20 µm.

**Figure 4 – figure supplement 1.**
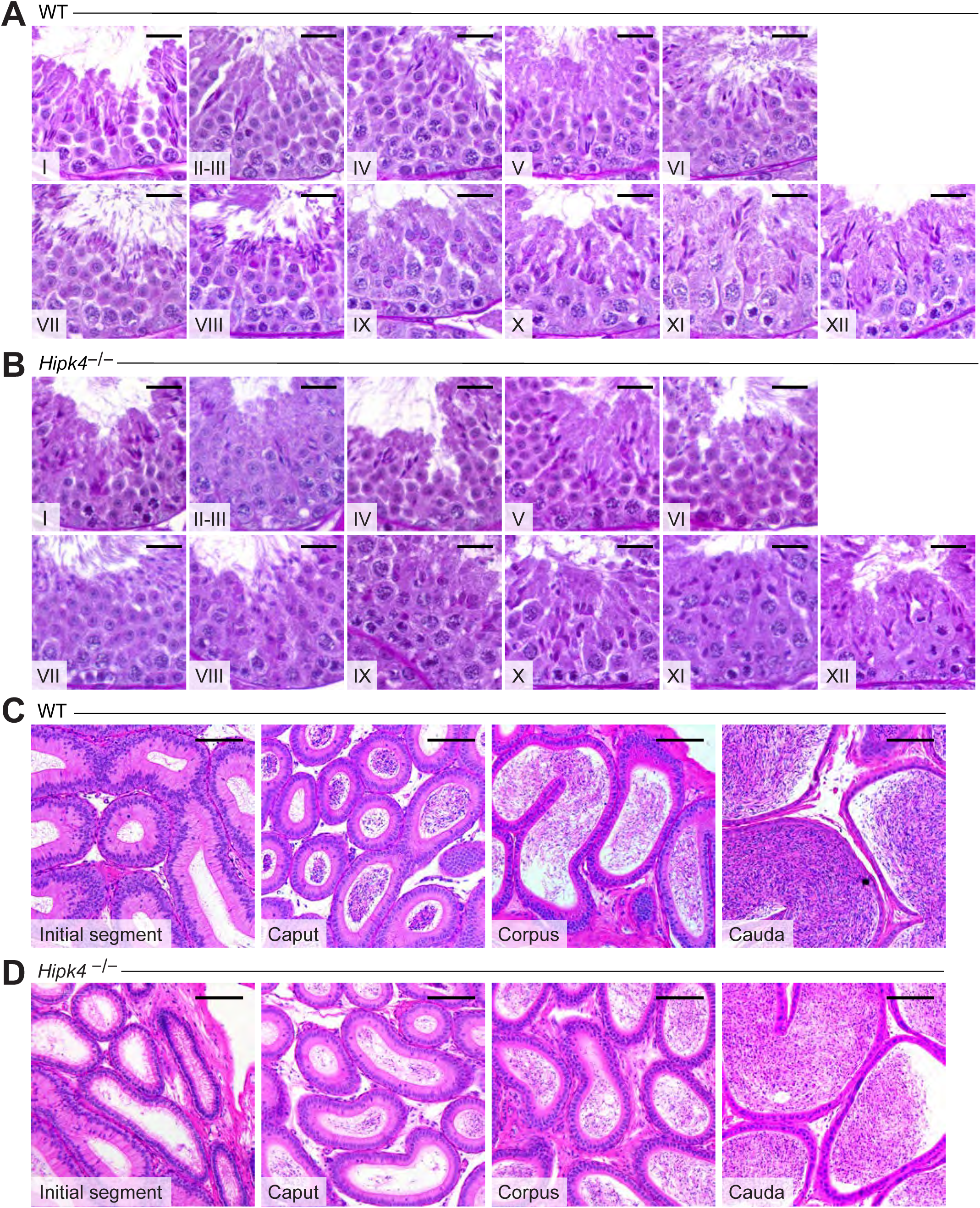
Comparison of the seminiferous epithelium and epididymis of WT and *Hipk4* knockout mice. (A-B) Fixed, paraffin-embedded testis sections from WT (A) and *Hipk4*^−/−^ (B) mice stained with PAS reagents. Stages for the tubules shown in each micrograph are indicated. (C-D) Fixed, paraffin-embedded epididymis sections from WT (C) and *Hipk4*^−/−^ (D) mice stained with hematoxylin and eosin. Epididymal regions associated with each section are indicated. Scale bars: 20 µm.

**Figure 4 – figure supplement 2.**
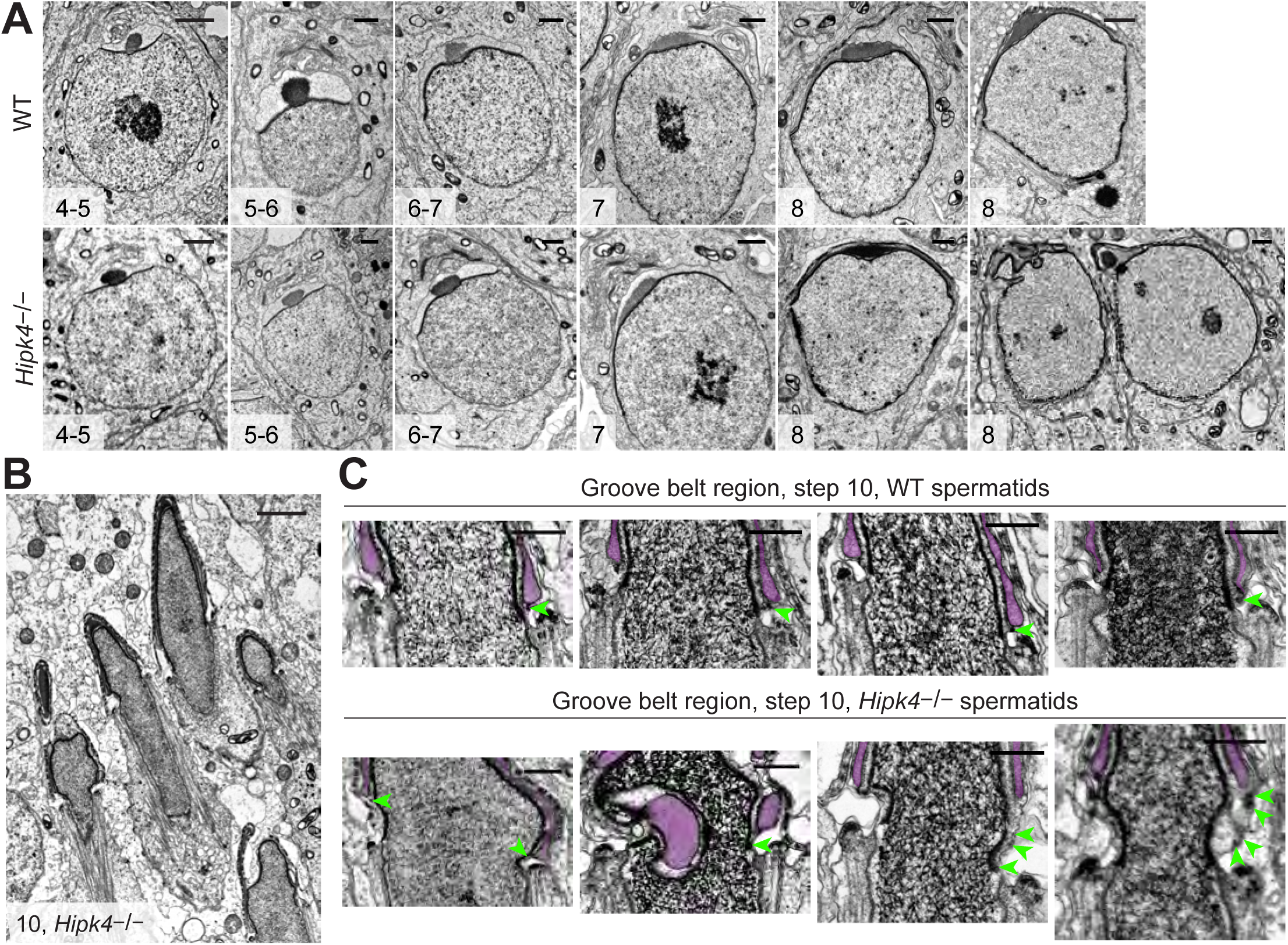
HIPK4 null spermatids exhibit acrosome-acroplaxome defects. (A) TEM images of step 4-8 spermatids from adult WT and *Hipk4^−/−^*mice. (B) TEM images of step 10 *Hipk4^−/−^*spermatids. (C) Higher magnification TEM images of the groove belt region in step 10 WT and *Hipk4^−/−^*spermatids. Acrosomes are false-colored purple, and green arrowheads label electron densities in the acroplaxome that normally are associated with the acrosome or plasma membrane. Scale bars: A, 1 µm; B, 2 µm; C, 1 µm.

**Figure 4 – figure supplement 3.**
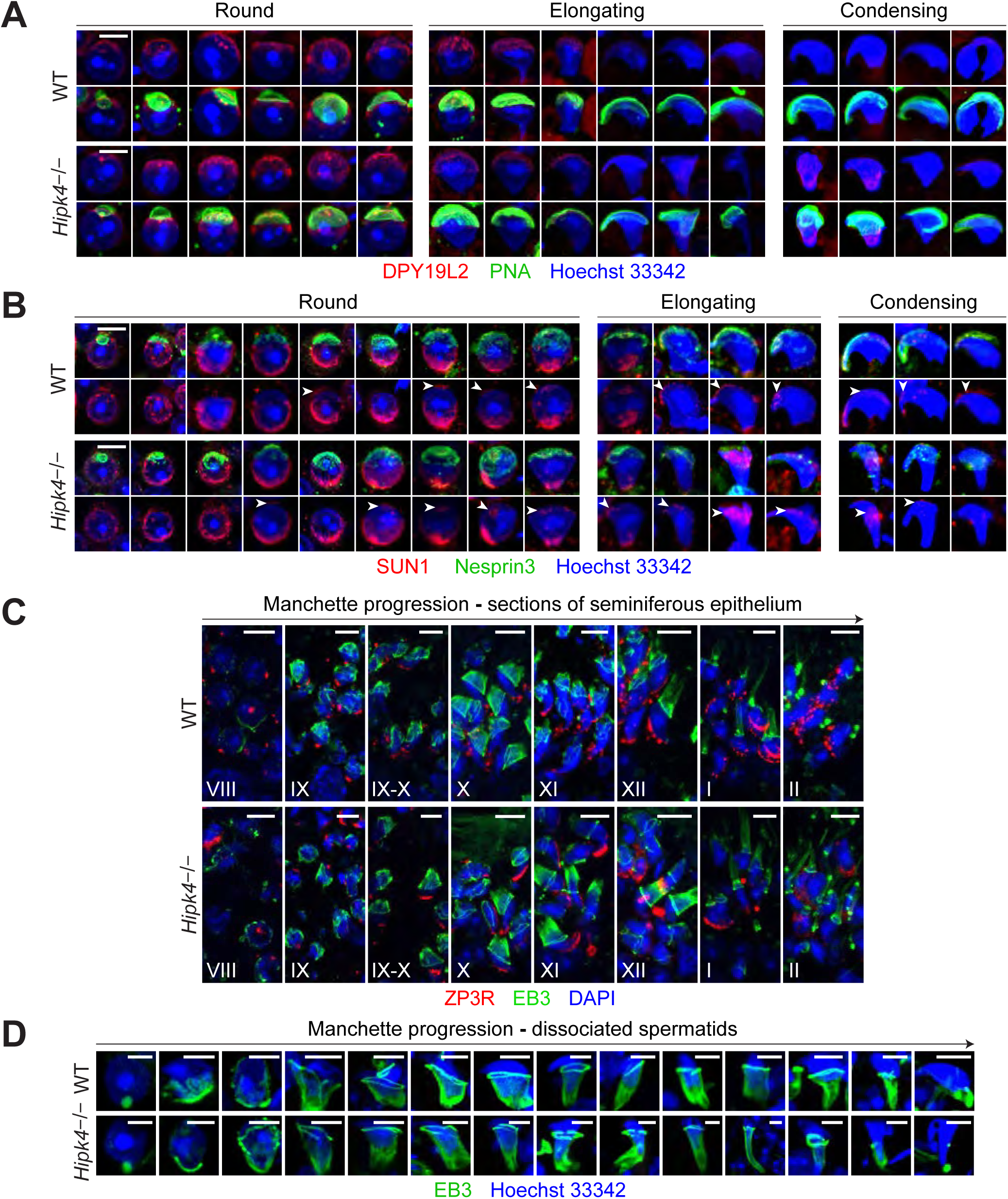
HIPK4 does not regulate the localization of anterior LINC complexes or manchette dynamics. (A) Immunofluorescence imaging of the inner nuclear membrane protein DPY19L2 (red) in dissociated spermatids. The acrosome is labeled with FITC-PNA (green), and nuclei are stained with Hoechst 33342 (blue). (B) Immunofluorescence imaging of the LINC complex proteins SUN1 (red) and nesprin3 (green) in dissociated spermatids. Nuclei are stained with Hoechst 33342 (blue). (C) Immunofluorescence imaging of EB3 (green) and ZP3R (red) in testis cryosections. Nuclei are stained with DAPI, and stages of each section are indicated. (D) Immunofluorescence imaging of EB3 (green) in dissociated spermatids. Nuclei are stained with Hoechst 33342 (blue). Scale bars: A-B (scale bars are representative for all images across the row), 2 µm; C, 5 µm; D, 2 µm.

